# Engineered Antigen-Binding Fragments for Enhanced Crystallisation of Antibody:Antigen Complexes

**DOI:** 10.1101/2023.07.06.548021

**Authors:** Heather Ann Bruce, Alexander U. Singer, Ekaterina Filippova, Levi Lynn Blazer, Jarrett J. Adams, Leonie Enderle, Moshe Ben-David, Elizabeth H Radley, Daniel YL Mao, Victor Pau, Stephen Orlicky, Frank Sicheri, Igor Kourinov, Shane Atwell, Anthony A. Kossiakoff, Sachdev S Sidhu

## Abstract

The atomic-resolution structural information that X-ray crystallography can provide on the binding interface between a Fab and its cognate antigen is highly valuable for understanding the mechanism of interaction. However, many Fab:antigen complexes are recalcitrant to crystallisation, making the endeavour a significant effort with no guarantee of success. Consequently, there have been significant steps taken to increase the likelihood of Fab:antigen complex crystallisation by altering the Fab framework. In this investigation, we applied the surface entropy reduction strategy coupled with phage-display technology to identify a set of surface substitutions that improve the propensity of a human Fab framework to crystallise. In addition, we showed that combining these surface substitutions with previously reported Crystal Kappa and elbow substitutions results in a striking improvement in Fab and Fab:antigen complex crystallisability, revealing a synergistic relationship between these sets of substitutions. Through comprehensive Fab and Fab:antigen complex crystallisation screenings followed by structure determination and analysis, we defined the roles that each of these substitutions play in facilitating crystallisation and how they complement each other in the process.

## INTRODUCTION

X-ray crystallography is a powerful means for determining structures of proteins and protein:protein interactions at atomic detail ^1,2^. This level of detail is particularly valuable for the study of antibody:antigen interactions, which can be used to characterise binding mechanisms, to identify determinants of affinity and specificity, and to optimize therapeutic antibodies ^3–5^. Structural characterization of full-length IgGs associated with antigen is immensely challenging due to the inherent flexibility of the IgG tertiary structure; in particular, the flexibility of the hinge regions connecting the antigen-binding fragments (Fabs) to the crystallisable fragment (Fc). To facilitate antibody:antigen complex crystallisation, IgGs are typically truncated to Fabs or even smaller single-chain variable fragments (scFvs) that contain the entire antigen-binding site (i.e. paratope) but are less dynamic, and thus, enable more facile crystallization and structure elucidation for characterisation of the paratope:epitope interface. Fabs in particular are well-structured, β-sheet-rich, unmodified heterodimers of ideal size for crystallography [6] that can even be used to aid the process as chaperones ^7–9^. Nevertheless, the generation of high-quality crystals from Fab:antigen complexes remains a bottleneck for structural studies, as many complexes are recalcitrant to crystallisation.

Crystallisation involves the formation of a lattice structure supported by stable and regular anisotropic packing interactions between neighbouring particles ^10,11^. Certain energy barriers must be overcome for this arrangement to occur - such as the entropic cost from the loss of translational and rotational degrees of particle freedom, and a reduction in local or overall conformational flexibility ^10,12^. Penalties to the Gibbs free energy of crystallisation must be offset by those that favour the process, such as an enthalpic loss from bond formation and an entropic gain upon solvent expulsion from crystal contact sites ^10,12^. It is therefore highly desirable to reduce the entropic cost of lattice formation by reducing unnecessary high-entropy elements from the protein structure before crystallisation. The surface entropy reduction (SER) strategy can provide an effective means for enabling crystallisation and generating superior diffraction-quality crystals in cases where the wild-type protein does not crystallise or provides poor quality crystals ^13–15^. In principle, the SER process involves changing the composition of amino acids on the protein surface which are entropically less favourable for mediating crystal lattice contacts ^16,17^. In practice, surface residues with relatively high side chain conformational entropy (e.g. Lys, Arg, Asp, Glu) are substituted with smaller amino acids (e.g. Ala, Gly, Ser) by site-directed mutagenesis ^14,16,18^. The resulting SER-modified sites are anticipated to enable sampling of different crystal lattice packing arrangements by providing additional or alternative points of contact to those in the WT protein [15].

Crystallisation can also be enhanced by the deletion of flexible regions by site-directed mutagenesis or limited proteolysis *in situ* [17]. Given the immense number of characterized natural and synthetic antibodies, improved methods for crystallisation of antibody:antigen complexes could have broad impact across basic research and drug development. Optimization of Fab entropy without compromising fold and function could systemically improve structural studies of these highly conserved proteins and their complexes. Indeed, improvement in the diffraction resolution of several Fab:antigen complexes was achieved by making substitutions in the heavy “elbow” region (which connects the constant and variable domains of the heavy chain in the Fab framework), resulting in a reduced Fab elbow angle range of 164-186° ^19,20^. The altered conformational state that the elbow ‘switch ‘substitution confers to the Fab framework was shown to result in fewer Fab molecules in the crystallographic asymmetric unit, thus making the structures easier to solve and refine [19].

Another approach to improving Fab crystallisation was recently reported [21]. Using the wealth of Fab apo form and Fab:antigen complex crystal structures available in the Protein Data Bank (PDB), a comprehensive analysis of Fab-mediated crystallisation packing sites was performed. This analysis revealed that a subset of rabbit Fab crystal structures contained a β-sheet stacking interface between the light and heavy chain constant domains of neighbouring Fab molecules in the crystal lattice. To promote this packing mechanism in human Fabs, the authors grafted the rabbit-derived structural segment from the light chain, which they termed “Crystal Kappa”, onto the corresponding region in a human Fab framework, which resulted in the expected crystal lattice packing [21].

In this study, we have employed the SER strategy in combination with phage display technology to identify a small group of surface substitutions that provide a human Fab framework with improved crystallisability. We then performed a comprehensive analysis of the crystallisability of Fabs containing these substitutions alone or in combination with previously reported Crystal Kappa and elbow substitutions ^19,21^ to identify highly crystallisable Fab proteins. Consequently, we have developed a crystallisation platform for facile and efficient crystallisation of Fab:antigen complexes using this optimized system.

## RESULTS

### Selection and characterisation of Fab variants with reduced surface entropy

We aimed to further improve the crystallisability of the Crystal Kappa Fab framework [21] by identifying positions in the light chain constant (CL) domain, which may be amenable to the SER strategy [17]. We compiled a panel of 43 Fab and Fab:antigen complex crystal structures with a common human framework that we and others have used for antibody humanization and generation of synthetic antibodies by phage display (**Supplementary Table S1**). The structures were processed by the CryCo (<u>Cry</u>stal <u>Co</u>ntact) server [22] to evaluate the relative degree to which each surface residue participates in crystal lattice packing interactions. We identified six surface residues in the CL domain with long side chains and >30% crystal contact binding recurrence and modelled them on a human Fab framework (contact residues, **Figure 1A**). Additionally, we identified twelve surface residues with long side chains neighbouring the contact residues (adjacent residues, **Figure 1A**).

**Figure 1.**
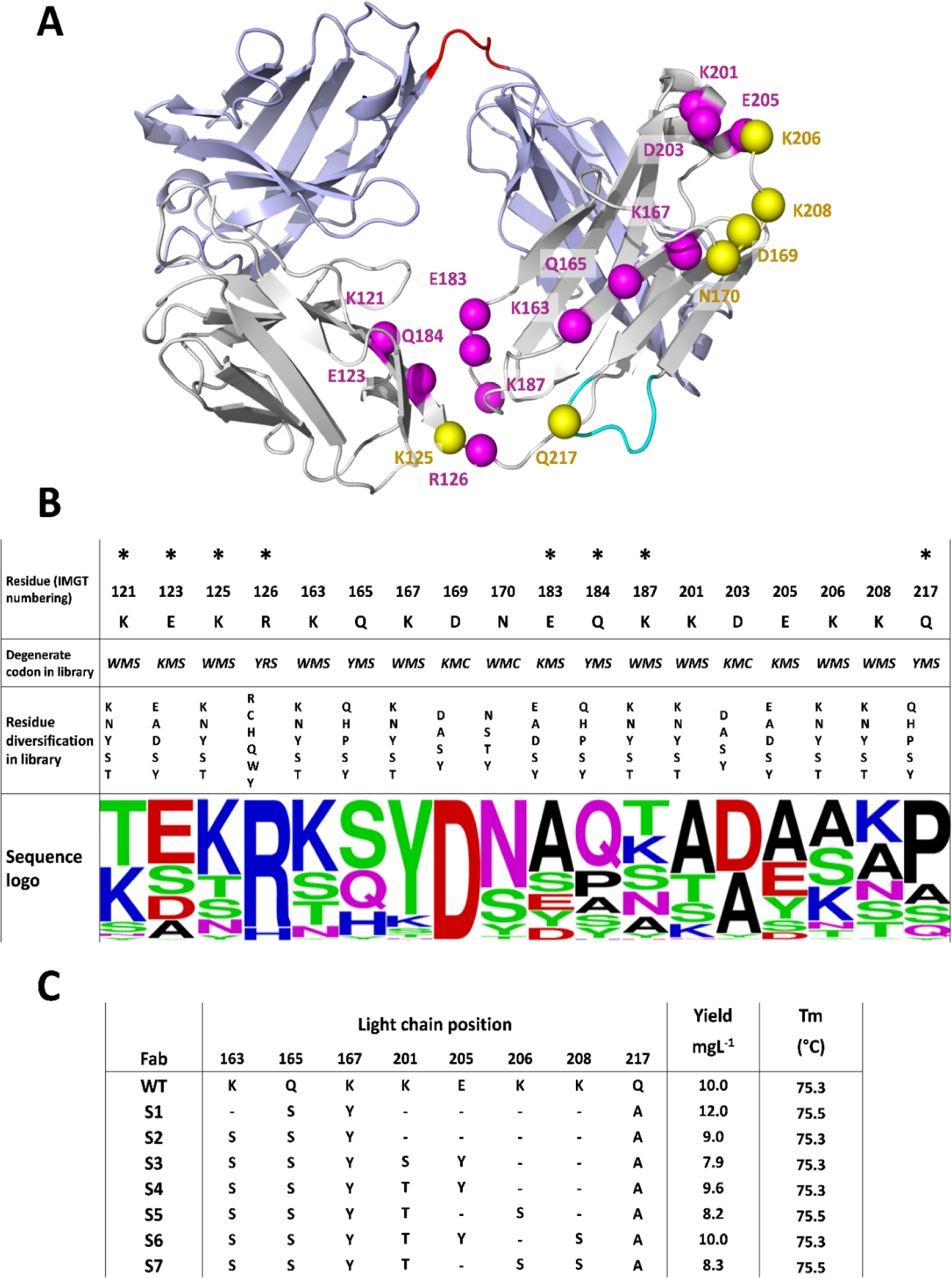
Fab variants with SER substitutions. **(A)** The library design mapped onto the Fab structure. The Fab framework mainchains are shown as ribbons (PDB accession code: 3L95). The light chain is colored grey, except residues in the Crystal Kappa graft, which are colored cyan. The heavy chain is colored light blue, except the elbow region, which is colored red. Positions that were randomized in the phage-displayed libraries are labelled with IMGT numbering, which is used throughout [25], and are shown as spheres, colored yellow or magenta for contact or adjacent residues, respectively (see main text for further details). **(B)** Library design and selection results. The WT sequence is shown at the top. Each position was diversified in one of two libraries, and those in the first library are indicated by asterisks (*). The degenerate codon used at each position is shown below the WT sequence in *italics* (W = A/T, M = A/C, S = G/C, Y = U/C, K = G/T and R = A/G), and the amino acids encoded are listed below. The sequence logo below each position depicts the prevalence of amino acids amongst clones selected for binding to the EPHA2 ECD, the cognate antigen for Fab F1. **(C)** Fab variants chosen for characterization in crystal screens. The WT Fab sequence is shown at the top, and the sequences of the seven Fab variants (S1-7) are shown below. Dashes indicate identity with the WT sequence. The yield of protein from 1-L of bacterial culture and the T_m_ determined by differential scanning fluorimetry are shown to the right.

To explore alternative lower entropy residues at the contact and adjacent positions in a rapid manner, we designed two phage-displayed libraries, that together, targeted all eighteen residues. At each position, we used a degenerate codon that encoded the WT residue and several alternative residues with smaller side chains (**Figure 1B**). The libraries were constructed in the context of a Fab (named F1) that bound to the C-terminal fibronectin domain (FN2) of the human receptor tyrosine kinase (RTK) EPHA2 and were subjected to binding selections for the extracellular domain (ECD) of EPHA2 [23] (Adams et al.,– *manuscript in preparation*). Positive binding clones were subjected to DNA sequence analysis and unique protein sequences were aligned to determine the prevalence of each allowed amino acid at each position, which was depicted as a sequence logo (**Figure 1B**). Aside from D169 which was conserved as the WT, diverse sequences were observed at the targeted positions, consistent with most surface residues not contributing significantly to the stability of the protein fold.

Numerous variants representing diverse sets of substitutions were recombinantly expressed and purified from *Escherichia coli* and screened for yield and thermostability. The screen identified two Fab F1 variants (Fab^S1^ and Fab^S2^) with yields and melting temperatures (T_m_) very similar to Fab^WT^ F1. Both variants shared three common substitutions (Q165S, K167Y, Q217A), with Fab^S2^ containing a fourth substitution (K163S). Based on the prevalence of substitutions at other positions, we constructed a panel of 27 additional variants containing two, three or four substitutions, in addition to the four substitutions found in Fab^S2^ (**Supplementary Figure S1**). The panel was screened for yield and thermostability, and this process yielded five additional candidates with yields and thermostabilities comparable to Fab^WT^ F1 (**Figure 1C**). Thus, the selection and screening process resulted in a panel of seven Fab variants containing three to seven substitutions of highly flexible surface side chains with less flexible side chains, but with yields and thermostabilities comparable to Fab^WT^ F1. Moreover, these Fab F1 SER-variants exhibited no evidence of degradation, oligomerisation, or aggregation detected by size exclusion chromatography (SEC) and native gel electrophoresis (**Supplementary Figure S2**).

### Crystallisation screens of Fab F1 variants with reduced surface entropy

With each of the seven Fab F1 SER-variants containing substitutions conferring reduced surface entropy (**Figure 1C**), extensive crystallisation screening was undertaken to gauge the relative degree of crystallisability compared with Fab^WT^ F1 (**Figure 2A**). For each, we screened 576 conditions composed of six 96-well plates, with each plate containing one of the following: sparse matrix screens JCSG-plus HT-96 Eco (Molecular Dimensions) and INDEX HT (Hampton Research), salt screen SaltRX HT (Hampton Research), pH screen with a varying PEG/ion environment PACT Premiere HT-96 (Molecular Dimensions), and two screens favorable for testing monoclonal antibodies; GRAS Screen^TM^ 1 and GRAS Screen^TM^ 2 (Hampton Research). Any condition which yielded crystalline material, irrespective of morphology or size of crystal, was considered a single crystal hit [11]. The INDEX screen was the most effective as it generated multiple crystal hits with all Fabs. All SER-variants provided fewer crystal hits than Fab^WT^, except Fab^S1^, which contained only three substitutions (Q165S, K167Y and Q217A) but provided the most hits, and also, exhibited higher protein yield and thermostability than Fab^WT^ F1 (**Figure 1C**). Consequently, the S1 substitutions were taken forward for further analysis and optimization.

**Figure 2.**
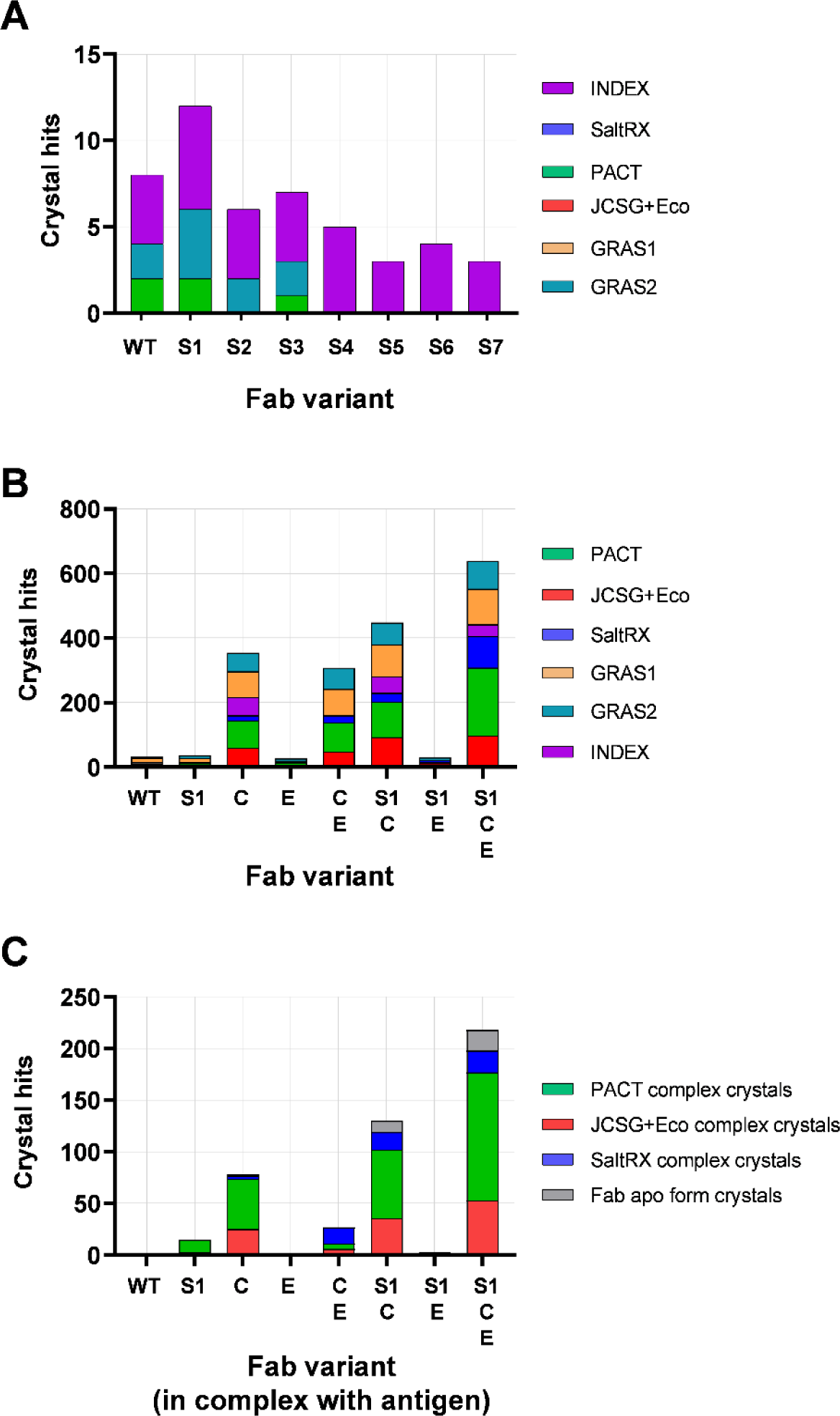
Results of crystal screens for Fab variants. **(A)** Results for WT and SER variants (S1-S7) of Fab F1. The number of crystal hits (y-axis) obtained for each Fab variant (x-axis) are shown after screening a total of 576 conditions (96 conditions for each of the six indicated screens). **(B)** Aggregate results for WT and variants of Fabs F1, 14386, and F4.A containing S1, Crystal Kappa (C), elbow (E), or combination substitutions. The number of crystal hits (y-axis) obtained for each set of three Fab variants (x-axis) are shown after screening a total of 1,728 conditions (96 conditions for each of the six indicated screens for each of the three Fabs) (See **Supplementary Figure S4** for individual Fab results). **(C)** Aggregate results for WT or indicated variants of Fab F1 in complex with EPHA2-FN2, and the dual-antigen-binding Fab 14386 in complex with antigen-A or -B. The number of crystal hits (y-axis) obtained for each set of three Fab:antigen complexes with the indicated Fab framework (x-axis) are shown after screening a total of 864 conditions (96 conditions for each of three indicated screens for each of the three Fab:antigen complexes). (See **Supplementary Figure S6** for individual Fab:antigen complex results).

### Crystallisation screens of Fab variants combining S1, Crystal Kappa, and heavy-chain elbow substitutions

We next compared the crystallisation of Fab^WT^ with Fab variants containing S1 substitutions (Fab^S1^), Crystal Kappa substitutions (Fab^C^) [21], heavy-chain elbow substitutions (Fab^E^) [19], pairwise combinations of substitutions (Fab^S1C^, Fab^S1E^, Fab^CE^), or all three sets of substitutions (Fab^S1CE^). When combining the S1 and Crystal Kappa substitutions, we used the Q165S and K167Y substitutions from Fab^S1^, but because position 217 differs between the two, we used the Q217G substitution from Crystal Kappa rather than the Q217A mutation from Fab^S1^, for consistency with the original study [21]. In addition, as Fab complementarity determining regions (CDRs) may contribute to and influence crystal packing interactions, we screened three Fabs with identical frameworks but distinct CDRs and antigen recognition. Along with Fab F1, we screened Fab F4.A, which recognizes the N-terminal cysteine-rich domain (CRD) of Frizzled-4 (FZD4) (Blazer et al., – *manuscript in preparation*), and a 2-in-1 Fab (PMID-19299620) named 14836, which recognizes two distinct antigens separately.

Fab proteins were recombinantly expressed in mammalian Expi293 cells and were purified to homogeneity without any signs of degradation, oligomerisation, or aggregation detected by denaturing and native gel electrophoresis. Consistently higher yields were obtained for F1, F4.A and 14846 Fab variants containing the S1 substitutions (**Supplementary Figure S3A**). In addition, the S1 substitutions conferred a marginal increase to the T_m_ as assessed by differential scanning fluorimetry (tested for Fab F1 variants only) (**Supplementary Figure S3B**). For each of the three distinct paratopes, we subjected Fab^WT^ and the seven framework variants to crystallisation screens with the same 576 conditions described above for the panel of SER-Fab variants. The number of conditions that generated crystal hits for each of the 24 Fab framework/paratope combinations were determined and plotted separately (**Supplementary Figure S4**). We also plotted the aggregate of crystal hits for eight groups, with each group containing three Fabs with distinct paratopes but identical frameworks (**Figure 2B**). This analysis showed that frameworks that contained the Crystal Kappa substitution exhibited dramatically enhanced crystallisation compared with those that did not. Crystallisation was further enhanced by combining the S1 substitutions with the Crystal Kappa substitution, and in many conditions, the crystal morphology improved also, revealing beneficial complementarity between these sets of substitutions (**Supplementary EXCEL document**). Crystallisation was enhanced still further by including the elbow substitutions with the Crystal Kappa and S1 substitutions. Overall, the two best performing frameworks were Fab^S1C^ and Fab^S1CE^, for which an amazing 26% or 37% of conditions, respectively, generated crystal hits across an extensive range of screening conditions, compared to only 2% for the Fab^WT^ framework.

### Crystallisation screens of Fab variants in complex with antigens

Next, we screened the Fab variants of F1 in complex with EPHA2-FN2, and the Fab variants of 14836 in complex with either antigen-A or antigen-B. For Fab^WT^ and each of the seven framework variants, the complex was formed by mixing Fab with antigen at a 1:2 molar ratio and a final concentration of 7 mg/ml. Each set of the Fab:antigen combinations was subjected to the three crystallisation screens that yielded the most hits for the apo Fab screens (JCSG+Eco, PACT, and SaltRX), and thus, a total of 2,304 conditions were screened for each antigen (288 conditions for each of the eight Fab frameworks). Apo Fab drops were set up in parallel to evaluate whether apo Fab crystals may have formed in the complex drops by comparing crystal morphology (**Supplementary EXCEL document**). In addition, for the Fab-14836:antigen-A complex screens, crystal composition was verified by picking crystals directly from the 96-well drops when possible and, after washing away precipitant, analysing crystal composition by denaturing polyacrylamide gel electrophoresis (**Supplementary Figure S5**). As would be expected, a small number of conditions generated putative apo Fab crystals in the complex crystallisation drops (**Figure 2C, grey bars**).

For the Fab-F1:EPHA2-FN2 complex screens, the most hits were obtained with Fab^C^, many hits were also obtained with Fab^S1^, Fab^CE^, Fab^S1C^ and Fab^S1CE^, one good hit was obtained with Fab^S1E^, but no hits were obtained with Fab^WT^ or Fab^E^ (**Supplementary Figure S6A**). For the Fab-14836:antigen-A complex screens, the most hits were obtained with Fab^S1CE^, many hits were also obtained with Fab^C^, Fab^S1C^, and Fab^CE^, a couple of hits were obtained with Fab^S1E^, but no hits were obtained with Fab^WT^, Fab^E^ or Fab^S1^ (**Supplementary Figure S6B**). For the Fab-14836:antigen-B complex screens, numerous hits were obtained with Fab^S1CE^ and Fab^S1C^, a couple of hits were obtained with Fab^C^ and Fab^CE^, but no hits were obtained with the other frameworks (**Supplementary Figure S6C**). Among the three sets of substitutions for the three Fab:antigen complexes in aggregate, the Crystal Kappa framework (Fab^C^) provided the greatest number of crystal hits, which were increased by combining Crystal Kappa with the S1 substitutions (Fab^S1C^) and even further increased with the addition of the elbow substitutions (Fab^S1CE^) (**Figure 2C**). Indeed, for the three complexes, the Fab^C^, Fab^S1C^, and Fab^S1CE^ frameworks generated hits in 9%, 14%, and 23% of conditions screened, respectively, in contrast to the Fab^WT^ framework which did not generate any hits at all.

### Structural analysis of Fab F1 with Fab^S1^, Fab^C^ and Fab^S1CE^ frameworks

To investigate how the S1, Crystal Kappa, and elbow substitutions – and their combinations -impact crystal lattice packing, we solved the crystal structures of Fab F1 with three distinct frameworks: Fab^S1^, Fab^C^, and Fab^S1CE^ (**Crystallography data Table**). In the crystallisation conditions which generated hits for both Fab^WT^ F1 and Fab^S1^ F1, crystals of the latter presented better morphology (i.e., possessed a smoother, non-striated surface) (**Supplementary EXCEL document**). A condition from the INDEX screen with ammonium sulfate and high molecular weight PEG was selected for crystallisation optimisation of both Fab^WT^ F1 and Fab^S1^ F1, resulting in plate-shaped crystals that diffracted to 4.2 Å or 3.5 Å, respectively. The quality of the Fab^WT^ F1 dataset was too poor to solve with confidence. However, the Fab^S1^ F1 structure was solved with an orthorhombic crystal system and P2_1_2_1_2_1_ symmetry. Within the asymmetric unit, Fab^S1^F1 molecules form a hexameric arrangement **(Figure 3Ai)**, led by aromatic residues in the exposed CDRs. The S1 substitutions Q165S and K167Y - along with D169, N170, A171, and L172 surface residues in the nearby loop region - were found to participate in several different crystal lattice packing interactions between neighbouring Fab hexamers in the Fab^S1^ F1 structure (**Supplementary Table S2**). In one of the Fab^S1^ F1 crystal lattice packing sites, hydrogen bond interactions can be observed between S1 substitutions Q165S/K167Y and residues T215*/Q216* from a neighbouring Fab CH domain (**Figure 3Bi**) (NB: asterisks are used throughout the main text and Figure legends to distinguish heavy-chain residues from light-chain residues). Meanwhile, in another Fab^S1^ F1 lattice packing site, K167Y forms a hydrophobic packing interaction against L172 in the nearby loop region of the same Fab molecule, which in turn packs against I219* from the CH domain in a neighbouring Fab molecule (**Figure 3B (ii)**). At a third Fab^S1^ F1 crystal lattice packing site, the K167Y substitution - together with N170 and A171 in the neighbouring loop region of the CL domain - facilitates an interaction through hydrogen bond and hydrophobic interactions with R95* and P15* residues of an adjacent Fab molecule VH domain (at a site distal to the CDRs) (**Figure 3B (iii)**). The myriad of different Q165S/K167Y-mediated crystal lattice packing interactions like these ones (and others observed in the Fab^S1^ F1 structure - see **Supplementary Table S2**), suggests that the S1 substitutions Q156S/K167Y provide a versatile crystal lattice packing site in the Fab CL domain.

**Figure 3.**
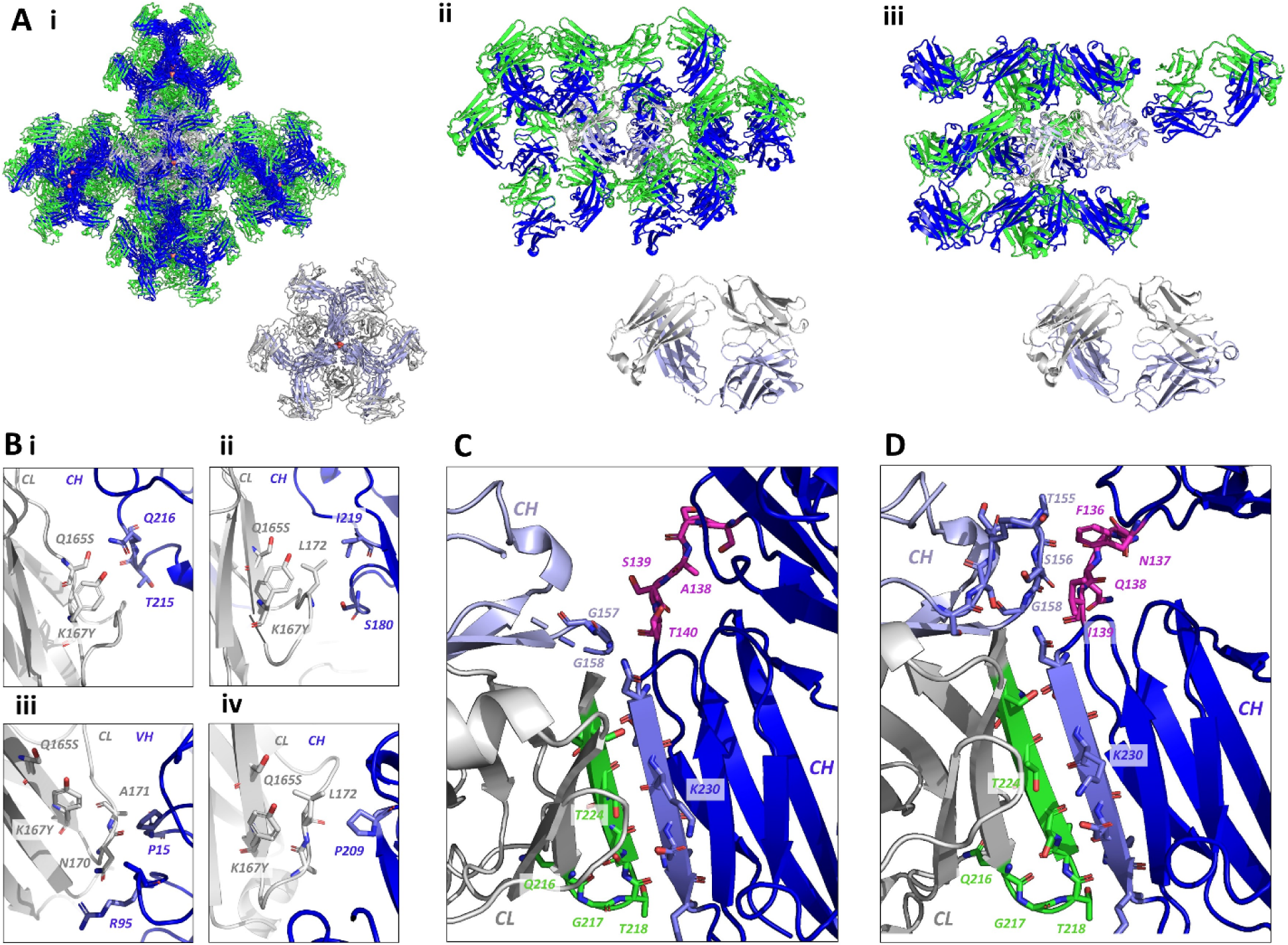
The impact of S1, Crystal Kappa and elbow substitutions on crystal lattice packing interactions of Fab F1. **(A)** Crystal lattice packing arrangement with symmetry mates (upper panel), and asymmetric unit (lower panel), of Fab F1 with the following frameworks: **(i)** Fab^S1^ (P2_1_2_1_2_1_ space group) (sulfate ions depicted as red spheres), **(ii)** Fab^C^ (C2 space group), and **(iii)** Fab^S1CE^ (P4_2_2_1_2 space group). The heavy and light chains of the asymmetric unit are colored light blue or grey, respectively. For other selected Fab particles that form part of the crystal lattice, heavy and light chains are colored dark blue or green, respectively. **(B)** Selected packing interactions mediated by the S1 substitutions Q165S/K167Y in **(i, ii and iii)** the Fab^S1^ F1 crystal lattice, and **(iv)** the Fab^S1CE^ F1 crystal lattice. **(C)** The β-sheet stacking interaction (green/blue) mediated by Crystal Kappa between adjacent Fab molecules in the crystal lattice of Fab^C^ F1. The nearby WT elbow residues are shown in magenta. **(D)** The contiguous packing site in the Fab^S1CE^ F1 crystal lattice, formed by the Crystal Kappa signature interaction, and interactions between the elbow region (F136*, N137*, Q138*, I139*) (magenta) and residues in the packing Fab molecule CH domain (T155*, S156*, G158*). NB: asterisks are used throughout the main text and Figure legends to distinguish heavy-chain residues from light-chain residues.

Meanwhile, Fab^C^ F1 crystals were obtained in a high molecular weight PEG and low pH condition optimized from the broad screens, from which a 1.95 Å dataset was collected (**Crystallography data Table**). In the Fab^C^ F1 crystal structure (solved in a C2 space group with a monoclinic crystal system) the Crystal Kappa substitution enables stacking between the final N-terminal β-sheet structural segments of the CL and CH domains of two neighbouring Fab molecules in the crystal lattice, as intended (**Figure 3C**) [21]. Moreover, the Crystal Kappa-directed packing interaction in the Fab^C^ F1 structure overcame the CDR packing interactions that dominated the Fab^S1^ F1 crystal form, helping to reduce the number of Fab molecules in the asymmetric unit from six to one (**Figure 3A (ii)**). Notably, residues Q165 and K167 participate in a crystal lattice packing interaction, along with residue N170 in the nearby loop region (NB: N170 was identified in the CryCo analysis as having a high frequency of crystal lattice participation in apo Fab and Fab:antigen complex crystal structures – see **Figure 1 A**) (**Supplementary Table 2**).

As mentioned previously, in the broad screening we found that the Fab^S1CE^ framework facilitated crystallisation of Fab F1 significantly better than the Fab^S1C^ and Fab^CE^ frameworks, based on number of conditions which generated hits (152, 53, and 4 conditions generated hits for Fab^S1CE^, Fab^S1C^, or Fab^CE^, respectively), revealing striking complementarity between the three sets of substitutions (**Supplementary Excel document** and **Supplementary Figure S4A**). Moreover, the Fab^S1CE^ framework produced good-sized, non-striated Fab^S1CE^ F1 crystals, directly in the broad crystallisation screens, one of which diffracted to 2.6 Å (**Crystallography data Table**). The Fab^S1CE^ F1 crystal structure was solved in a higher symmetry space group (P4_2_2_1_2; tetragonal crystal system) than Fab^C^ F1, but with the signature packing interaction of Crystal Kappa preserved. In one of the crystal lattice packing sites observed in the Fab^S1CE^ F1 structure, a hydrophobic interaction is coordinated between the S1 substitution Q165S along with L172 in the principal Fab CL domain, and P209* in the CH domain of a neighbouring Fab molecule. Notably, the packing interactions mediated by the S1 substitutions in the Fab^S1CE^ F1 crystal lattice are distinct from those observed in the Fab^S1^ F1 crystal lattice (**Figure 3B**, compare i, ii and iii, with iv). In addition, the incorporation of the elbow substitution changed the elbow angle of Fab^S1CE^ F1 relative to Fab^C^ F1, as expected and described previously [19] (**Table 1**).

**Table 1.** Structural information for Fab and Fab:antigen complexes. The Fab elbow angle is described and defined in ^19,20^.

**Figure 6.**
| Fab/Fab:antigen complex | Mutations | Crystal system | Space group | Number of Fab/s or Fab:antigen complex/s in ASU | ASU Fab/s elbow angle (°) | PDB accession code |
| --- | --- | --- | --- | --- | --- | --- |
| Fab F1 | S1 | Orthorhombic | $P2_12_12_1$ | 6 | 185.2, 186.1, 184.9, 184.5, 184.6, 185.1 | 8T7F |
| | C | Monoclinic | $C2$ | 1 | 144.6 | 8T7G |
| | S1CE | Tetragonal | $P4_22_12$ | 1 | 171.8 | 8T7I |
| Fab-F1:EPHA2-FN2 complex | C | Monoclinic | $P2_1$ | 4 | 168.9, 168.4, 172.7, 172.7 | 8T9B |
| Fab-V1:VHH complex | - | Monoclinic | $C2$ | 9 | 173.8, 175.1, 172.3, 173.2, 185.3, 172.0, 170.6, 174.7, 173.4 | 7RTH<br>[24] |
| | E | Monoclinic | $P2_1$ | 2 | 172.6, 171.7 | 8T58 |
| | C | Orthorhombic | $P2_12_12_1$ | 1 | 143.5 | 8T6I |
| | CE | Tetragonal | $P4_32_12$ | 1 | 176.3 | 8T9Y |
| | S1CE | Orthorhombic | $P2_12_12_1$ | 1 | 165.3 | 8T8I |

The Fab^S1CE^ F1 crystal lattice arrangement also revealed that the proximity of the Crystal Kappa packing site and junction at the elbow substitution region results in the formation of a contiguous packing site between two Fab molecules in the crystal lattice (**Figure 3D**). Whilst the Crystal Kappa region facilitates packing with an adjacent Fab molecule through its signature β-sheet stacking interaction, residues from the elbow substitution region (F136*, N137*, Q138*, I139*) concomitantly form hydrogen bond and Van der Waals interactions with residues in the packing Fab CH domain (T155*, S156*, G157*), which presumably further stabilises the Fab:Fab packing arrangement induced by Crystal Kappa. Interestingly, the packing interaction site at the elbow junction is different in the Fab^C^ F1 and Fab^S1CE^ F1 crystal lattices (compare **Figure 3C** with **3D**). In contrast to the Fab^S1CE^ F1 crystal structure, just one residue (G157*) in the Fab^C^ F1 crystal structure forms an interaction with the WT elbow region (residue T140*) (**Supplementary Table S2**). Moreover, a significant portion of the loop region preceding residue S156* remains unresolved from the electron density in the Fab^C^ F1 structure, in contrast to the Fab^S1CE^ F1 structure in which the loop region is entirely resolved.

### Crystal lattice packing of Fab^C^ F1 in complex with the FN2 domain of EPHA2

To demonstrate the utility of the engineered Fab frameworks as crystallisation chaperones, we attempted to solve the structure of Fab F1 in complex with the FN2 domain of EPHA2 (EPHA2-FN2) using various Fab frameworks. In contrast to the Fab^WT^ framework, which did not yield any Fab^WT^-F1:EPHA2-FN2 complex crystals in the broad crystallisation screening, the Fab^S1^ framework generated rod-shaped complex crystals in many conditions in the JCSG+Eco and PACT screens, including some presenting good three-dimensional shape, whilst the Fab^S1E^ framework facilitated the formation of triangular prism-shaped complex crystals in a JCSG+Eco condition with optimisation potential (**Supplementary EXCEL document**). Meanwhile, the Fab^C^ and Fab^S1C^ frameworks generated the greatest number of complex crystal hits across all crystallisation screens tested, with the most promising morphology observed in a condition with a high concentration of sodium formate at high pH, which yielded thin, smooth, mica-like hexagonal plates. After crystallization optimization screening, the Fab^S1C^-F1:EPHA2-FN2 complex crystallised within a few days and produced a shower of tiny plate-shaped crystals. However, using the Fab^C^ framework, larger plate-like crystals formed slowly over more than a month, and these appeared best-suited for diffraction studies.

The highest resolution data collected for a Fab^C^-F1:EPHA2-FN2 crystal was fairly poor at 4.2 Å, with an early indication of a high solvent content (74%) from the Matthews coefficient calculations. Nevertheless, the structure was solved successfully by molecular replacement, with a monoclinic crystal system and a P2_1_ space group (**Crystallography data Table**) and represents the first structure of the FN2 domain of an EPH receptor bound to an antibody and not in the context of the full ectodomain. An analysis of the paratope:epitope interaction is presented elsewhere (Adams et al.,– *manuscript in preparation*). Here, we focus on the contribution that the Crystal Kappa substitution makes to the crystal packing in the Fab^C^-F1:EPHA2-FN2 complex structure, in the absence of the S1 and elbow substitutions. The asymmetric unit in the Fab^C^-F1:EPHA2-FN2 complex structure contains four Fab:antigen complexes assembled in an X-shaped arrangement led by antigen:antigen packing interactions at the center, which results in an outward splaying of the Fab molecules and large solvent cavities in the crystal lattice, thus explaining the high solvent content (**Figure 4A and B**). The Crystal Kappa signature packing interaction forms the principal crystallographic contact between neighboring asymmetric units (**Figure 4C**) (**Supplementary Table S3**). Moreover, the packing mediated by Crystal Kappa is consistent with that observed in the Fab^C^ F1 structure (**Figure 3C**), and that shown in the original study [21]. Meanwhile, the Fab CL residues Q165 and K167 (which differ in the S1 substitution) do not participate in any packing interactions.

**Figure 4.**
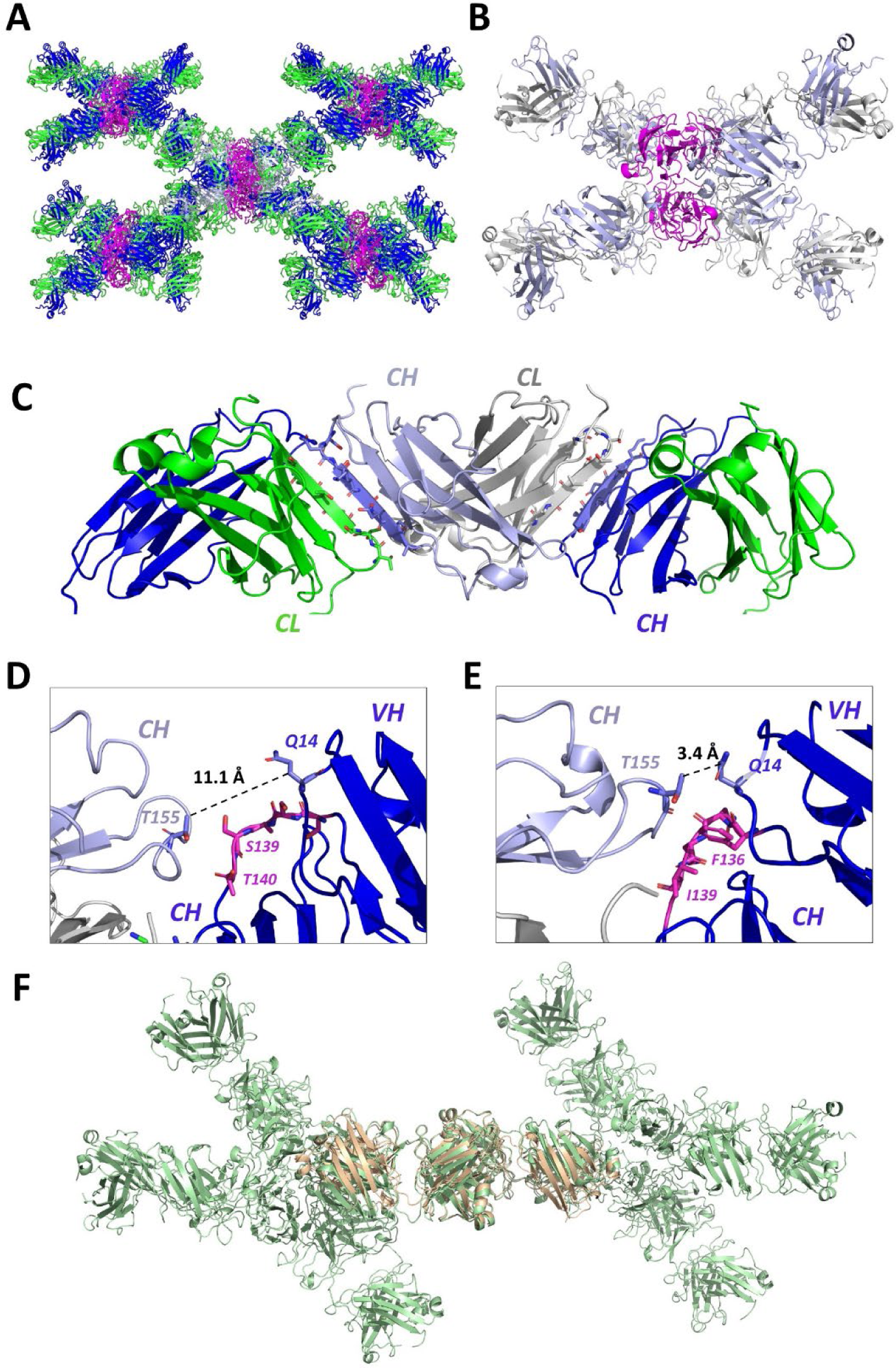
Analysis of Fab^C^-F1:-EPHA2-FN2 crystal lattice packing interactions mediated by Crystal Kappa. **(A)** Crystal lattice packing arrangement in the Fab^C^-F1:EPHA2-FN2 complex structure. A single asymmetric unit containing four Fab:antigen complexes (heavy and light chains colored light blue or grey, respectively) is shown with symmetry mates (heavy and light chains colored dark blue or green, respectively, and EPHA2-FN2 colored magenta). **(B)** The Fab^C^-F1:EPHA2-FN2 asymmetric unit. Four Fab:antigen complexes are connected by a tetrameric arrangement of the antigen at the centre, resulting in an X-shaped assembly with the four Fab molecules splayed outward. **(C)** Crystal Kappa signature β-sheet stacking, which forms the principal crystallographic packing interaction between neighbouring Fab light- and heavy-chain constant domains in the lattice structure. **(D and E)** Junction at the heavy-chain elbow region (pink) in **(D)** the Fab^C^-F1:EPHA2-FN2 complex structure, and **(E)** the apo Fab^S1CE^ F1 structure. Residues in the WT elbow region (S136*, S137*, A138*, S139*, T140*) do not form a packing interaction with the adjacent Fab CH domain (light blue) in the Fab^C^-F1:EPHA2-FN2 structure. In contrast, residues in the elbow substitution (F136*, N137*, Q138*, I140*) in the apo Fab^S1CE^ F1 structure, along with other residues in the VH domain (e.g. Q14*) form hydrogen bond and Van der Waals contacts with residues in the packing Fab CH domain (light blue). **(F)** Two Fab^C^-F1:EPHA2-FN2 ASUs which pack together in the crystal lattice are shown (pale green), with two Fab molecules from the apo Fab^S1CE^ F1 structure (beige) superimposed at the Crystal Kappa packing site. The differences in packing interactions at the elbow junction sites in the two structures results in a shift to the relative positions of the Fab variable domains, despite the similar elbow angle (see Table 1).

Despite the absence of the elbow substitution in the Fab^C^ framework, the four Fab molecules in the Fab^C^-F1:EPHA2-FN2 ASU have elbow angles similar to that exhibited by the apo Fab^S1CE^ F1 (**Table 1**), and within the expected range (164-186°) for crystal structures of Fabs that contain the elbow substitution [19]. Therefore, we might expect that combining the Crystal Kappa and elbow substitutions would facilitate crystallisation of the Fab-F1:EPHA2-FN2 complex just as well as the Crystal Kappa substitution alone. However, the Fab^CE^ and Fab^S1CE^ frameworks did not facilitate crystallisation of the Fab-F1:EPHA2-FN2 complex in the crystallisation screening in as many conditions, or even mostly in the same conditions, as the Fab^C^ framework (**Supplementary EXCEL Table** and **Supplementary Figure S6A**). As described in the previous section, residues in the elbow substitution region in the apo Fab^S1CE^ F1 structure form crystal lattice packing interactions with residues in a neighbouring Fab CH domain, as a result of the Crystal Kappa packing interaction located nearby (**Figure 3D**). The same interactions at the elbow junction are not preserved in the Fab^C^-F1:EPHA2-FN2 complex structure: not only is the distance between the elbow region and packing Fab CH domain too far (>4.3 Å) to confidently assign molecular interactions to, but the relative position of the packing Fab molecules is significantly altered (highlighted by comparing the distance between Q14* of the principle Fab VH domain and T155* of the adjacent packing Fab CH domain in the structures) (compare **Figure 4D** and **Figure 4E**) (**Supplementary Tables S2 and S3**). The altered crystal lattice packing at the elbow junction in the two structures results in a different relative placement of the packing Fab molecules when they are superimposed at their respective Crystal Kappa packing regions (**Figure 4F**). It is possible that the restrictions imposed on the crystal lattice packing in the Fab^C^-F1:EPHA2-FN2 complex structure (i.e. the EPHA2-FN2 antigen-led tetrameric arrangement and the Crystal Kappa-mediated stacking) make the lack of interactions between packing Fab molecules at the elbow junction more beneficial for crystallisation of this system. In this way, Fab-F1:EPHA2-FN2 asymmetric units can pack together with more flexibility and less rigidity, which makes the Fab^C^ framework more favourable than Fab^CE^ and Fab^S1CE^ frameworks for facilitating crystallisation of the Fab-F1:EPHA2-FN2 complex in this particular crystal lattice arrangement.

### Structural analysis of Fab-V1:VHH complexes with Fab^C^, Fab^E^, Fab^CE^, or Fab^S1CE^ frameworks

To further demonstrate the general and modular nature of the S1, Crystal Kappa and elbow engineered Fab frameworks, we selected another Fab:antigen complex for X-ray crystallography studies. Fab V1 (also known as NabFab) is a synthetic Fab with the same human framework as Fab F1 described in the previous sections, which binds to a conserved region in the framework of an autonomous VHH domain [24]. As Fab V1 binds distal to the VHH paratope, it does not interfere with the interactions of the VHH domain with its cognate antigen. Consequently, Fab V1 has been used as a fiducial marker and size enhancer in cryogenic electron microscopy studies of membrane proteins in complex with an existing VHH domain binding partner [24].

The Fab^WT^ V1 framework initially failed to produce suitable Fab-V1:VHH complex crystals for data collection, and crystals were only obtained after extensive optimization with crystallisation seeding [24]. We decided to investigate whether the crystallisability of the Fab-V1:VHH complex is enhanced when the S1, Crystal Kappa, and elbow substitutions are incorporated into the framework, thus improving the capacity of Fab V1 to act as a chaperone in the crystallisation of VHH:antigen complexes. To this end, Fab-V1:VHH complexes with various frameworks – Fab^C^, Fab^E^, Fab^CE^, Fab^S1E^ and Fab^S1CE^ - were prepared and screened alongside Fab^WT^ for comparison, using various JCSG+Eco, PACT, INDEX, ProComplex (NeXtal Biotechnologies) and PEG/ion HT^TM^ (Hampton Research) screens (specified in **Supplementary Figure S7**). For the Fab-V1:VHH complex, the Fab^E^, Fab^C^, Fab^CE^, and Fab^S1CE^ frameworks gave the highest number of hits. The Fab^S1E^ framework generated only a modest number of Fab-V1:VHH complex hits (comparable to Fab^WT^), but several good hits in terms of crystal morphology were obtained directly from the 96-well screens. Considering the number of successful conditions on the one hand, and crystal morphology on the other, it is clear that the engineered frameworks dramatically increased crystallisability of the Fab-V1:VHH complex compared with the Fab^WT^ framework.

Previously, the crystal structure of the Fab^WT^-V1:VHH complex was determined to a maximum resolution of 3.2 Å, in a low symmetry (C2) space group, containing nine complexes in the ASU [24] (PDB accession code: 7RTH) (**Figure 5A (i)**). In contrast, we were able to determine the crystal structure of the Fab-V1:VHH complex at good resolution using the Fab^E^, Fab^C^, Fab^CE^, and Fab^S1CE^ frameworks (ranging 2.2-2.5 Å), with all structures containing just one complex molecule in the ASU, except for the Fab^E^-V1:VHH complex crystals for which the ASU contained just two complexes (**Figure 5A (ii-v)**; **Crystallography data Table**). The Fab^C^ and Fab^S1CE^ frameworks yielded crystals with very similar crystal lattice packing: both structures were solved as orthorhombic crystal systems with P2_1_2_1_2_1_ space groups, but due to a difference in the Fab elbow angle, have a slightly different Matthew’s coefficient (Å^3^/Da) and solvent content (**Crystallography data Table** and **Table 1**). Meanwhile, the Fab^E^ and Fab^CE^ frameworks yielded crystals in monoclinic (P2_1_ space group) and tetragonal (P4_3_2_1_2 space group) crystal systems, respectively. Thus, our results reveal that crystal packing of Fab V1 in complex with the VHH domain has changed considerably by utilizing these alternative Fab frameworks, resulting in better diffracting crystals which required less optimization, and a reduction in the number of complex molecules in the ASU in addition to increased crystal lattice symmetry, making the structures easier to solve and refine.

**Figure 5.**
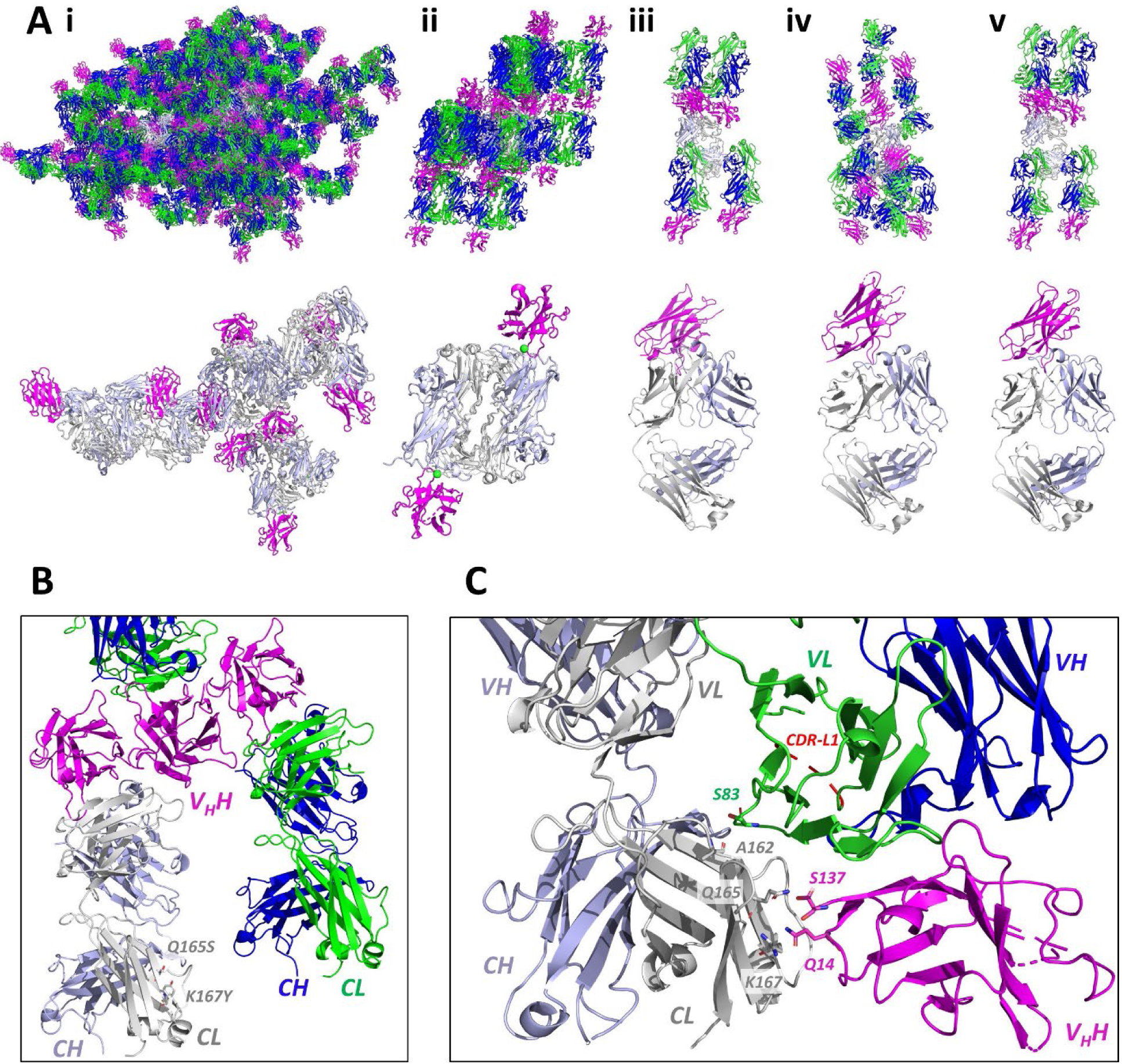
The impact of S1, Crystal Kappa and elbow substitutions on crystal lattice packing interactions of Fab-V1:VHH complexes. **(A)** Crystal lattice packing arrangement (upper panel) with symmetry mates, and asymmetric unit (lower panel), of Fab-V1:VHH complexes with the following frameworks: **(i)** Fab^WT^-V1:VHH (C2 space group) [24], **(ii)** Fab^E^-V1:VHH (P2_1_ space group) (Sodium ions depicted as green spheres), **(iii)** Fab^C^-V1:VHH (P2_1_2_1_2_1_ space group), **(iv)** Fab^CE^-V1:VHH (P4_3_2_1_2 space group), and **(v)** Fab^S1CE^-V1:VHH (P2_1_2_1_2_1_ space group). The Fab heavy- and light-chains in the asymmetric unit are colored light blue or grey, respectively, whilst the VHH domain is colored magenta. For the symmetry mates in the crystal lattice, heavy- and light-chains are colored dark blue or green, respectively. **(B)** The S1 substitution site (Q165S/K167Y) does not contribute to any crystal lattice packing interactions in the Fab^S1CE^-V1:VHH crystal structure. **(C)** Crystal lattice packing site in the Fab^CE^-V1:VHH structure involving the principle Fab CL domain (grey), and a neighbouring antigen and Fab VL domain (green). Fab CL residues Q165 and K167 form hydrogen bond interactions with Q14* and S137* in the packing VHH (antigen) molecule (coloured magenta). The packing is stabilised by hydrogen bond interactions between A162 of the Fab CL domain, and S83 of the packing Fab VL domain (amongst other interactions detailed in Supplementary Table S4). Due to the proximity of this packing site to the epitope:paratope interface in the Fab^CE^-V1:VHH crystal structure, there is a structural disruption to the CDR-L1 region that is left unresolved (red dotted loop), which does not occur in the CDR-L1 region in the Fab^S1CE^-V1:VHH crystal structure.

Notably, the crystal lattice packing arrangement in the Fab^S1CE^-V1:VHH structure is very different from that in the Fab^CE^-V1:VHH structure (**Figure 5A (compare iv and v)**). Inspection of the Fab^S1CE^-V1:VHH crystal lattice structure revealed that the S1 substitutions Q165S/K167Y do not mediate any packing interactions with neighbouring molecules (**Figure 5B**). In contrast, in the Fab^CE^-V1:VHH crystal lattice structure, residues Q165 and K167 form hydrogen bond interactions with residues Q14* and S137* in a nearby VHH molecule, stabilized through other interactions between residues in the CL domain of the principle Fab and residues of the packing Fab VL domain (**Figure 5C** and **Supplementary Table S4**). This results in higher symmetry and lower solvent content in the Fab^CE^-V1:VHH crystal lattice. However, the diffraction resolution was the same (2.5 Å) for both crystal structures, and the absence of any crystal lattice packing interactions so close to the paratope:epitope interface arguably makes the Fab^S1CE^ framework better than the Fab^CE^ framework for crystallization of this particular Fab:antigen complex. Indeed, the close packing between residues in the Fab CL domain and the adjacent Fab VL domain actually causes a disruption to the structural conformation of the CDR-L1 region in the Fab^CE^-V1:VHH complex structure (**Figure 5C**). Nevertheless, this intriguing result raises the possibility that, in some cases, the WT S1 site residues may provide a positive contribution to the crystal lattice packing. Taken together, the advantage that the S1 substitution confers to the crystallisability and crystal lattice packing is likely to be dependent on the Fab:antigen complex.

For all three Fab-V1:VHH structures containing the Crystal Kappa substitution (**5A (iii-v)**), its signature β-sheet-stacking interaction was present, highlighting its dominant role in the crystal lattice packing (e.g. see **Supplementary Figure S8** for Fab^C^-V1:VHH and Fab^S1CE^-V1:VHH Crystal Kappa packing interactions). Moreover, Fab-V1:VHH structures containing the Crystal Kappa substitution possess higher crystal lattice symmetry and have fewer molecules in the ASU than the Fab^WT^-V1:VHH and Fab^E^-V1:VHH complex structures (**Crystallography data Table**). In the Fab^C^-V1:VHH complex structure, the packing interactions at the elbow junction between residues in the elbow region and neighbouring Fab CH domain are limited, similar to that observed in the Fab^C^ F1 and Fab^C^-F1:EPHA2-FN2 structures (e.g. compare **Figure 3C** with **Supplementary Figure S8 (A)**). Meanwhile, as with the crystal packing observed at the elbow junction in the Fab^S1CE^ F1 structure, we found distances between residues in the elbow region and residues in the symmetry-related Fab CH domain suitable for packing interactions in the Fab^S1CE^-V1:VHH complex structure (compare **Figure 3D** with **Supplementary Figure S8 (B)**) (**Supplementary Table S4**).

## DISCUSSION

We utilised the SER strategy and phage display technology to identify substitutions that contribute to a favourable crystal lattice packing site on a Fab framework without destabilising the protein fold. The S1 substitutions (Q165S/K167Y/Q217A) increased Fab yield and thermostability, and enhanced crystallisation of both apo Fab F1 and the Fab-F1:EPHA2-FN2 complex, as evidenced by an increase in the number of crystal hits and the quality of crystal morphology. The Fab^S1^ F1 and Fab^S1CE^ F1 structures revealed that Q165S/K167Y form part of a crystal lattice packing site located in the Fab CL domain, with a myriad of different packing interactions observed. However, Q165S/K167Y are not involved in crystal lattice packing interactions in the Fab^S1CE^-V1:VHH complex structure. Thus, it appears that the S1 substitutions do not form a predictable crystal lattice packing site, but rather, they adapt to the requirements of individual Fab/Fab:antigen complex crystal lattice formations by acting as a versatile packing site sometimes in cooperation with nearby residues on the Fab surface.

Meanwhile, we found that the Crystal Kappa substitution provided a striking enhancement in the crystallisation of Fab and Fab:antigen complexes, in terms of number of crystal hits across a broad screen, and generally conferred an improvement in crystal morphology. Moreover, the Crystal Kappa signature β-sheet packing interaction was consistently observed in all structures analysed in this investigation and others, revealing its dominant contribution and influence on crystal lattice formation [21]. Furthermore, the Crystal Kappa substitution alone, and in combination with S1 and elbow substitutions, facilitated crystallisation of all four Fab:antigen complexes that we tested. Notwithstanding the excellent capacity of the Crystal Kappa substitution for enhancing crystallisability of the Fab framework, one caveat with using this substitution alone is that it could impose certain restrictions in crystal lattice packing, and when combined with a lack of other potential packing interactions, may result in a high solvent content and/or disruption of the native Fab:antigen complex. In such cases, it is advisable to use S1 and elbow substitutions, alone or in conjunction with the Crystal Kappa substitution, to capture alternative crystal lattice packing and symmetry forms. That being said, of the four Fab:antigen complexes that we tested, only Fab-F1:EPHA2-FN2 and Fab-V1:VHH complex crystals were generated by the S1 and elbow substitutions, respectively, revealing that when these substitutions are used independently, the crystallisation advantage that they confer to the Fab framework is highly dependent on the Fab:antigen complex.

The effect of the elbow substitution in reducing Fab conformational entropy by stabilising the elbow angle undoubtedly confers an advantage in the crystallisation of some Fab:antigen complexes [19]. Indeed, the elbow substitution was key to generating better diffracting Fab-V1:VHH complex crystals with higher crystal symmetry and fewer molecules in the ASU, as compared to the Fab^WT^ framework. However, the WT elbow may be considered a better alternative for some systems where the Fab structural conformation benefits from greater flexibility in order to adapt and accommodate the requirements of different Fab:antigen complexes in crystal lattice formation, particularly when combined with the Crystal Kappa substitution - as in case of the Fab-F1:EPHA2-FN2 complex.

Used independently, and in pair-wise combinations, the substitutions confer advantages to Fab:antigen complex crystallisation in distinct ways. However, a striking finding of this investigation was how well the three substitutions complement one another: the Fab^S1CE^ framework facilitated crystallisation of all Fab and Fab:antigen complexes tested, and it was the leading performer in terms of generating the most crystal hits with the best crystal morphology across the broad crystallisation screens. The Fab^S1CE^ F1 crystal structure revealed how the three substitutions can carry out their individual contributions to crystal lattice packing and complement one another: the S1 substitution forms a crystal lattice packing site, the Crystal Kappa concomitantly mediates its signature β-sheet stacking, while the elbow substitution reduces conformational flexibility by restricting the Fab elbow angle. Due to the proximity of the elbow region to the Crystal Kappa β-sheet stacking interaction, combining these two substitutions typically results in the formation of a contiguous packing site which probably contributes to the enhanced crystallisability of the Fab framework by stabilising and favouring the Crystal Kappa-mediated Fab:Fab packing mode in the crystal lattice.

In conclusion, the SER substitutions that we have generated can be combined with previously reported elbow [19] and Crystal Kappa [21] substitutions to provide a powerful toolkit for enhancing Fab and Fab:antigen crystallisation. For the best chance of Fab:antigen complex crystallisation success, we suggest using a combination of all three substitutions as a priority (Fab^S1CE^). Given an adequate supply of antigen protein, the Fab^S1C^ and Fab^C^ frameworks should also be utilised (**Figure 6**). In addition, with the aim to capture an alternative crystal lattice packing state, the Fab^E^ or Fab^S1E^ frameworks should also be tried.

**Figure 6.**
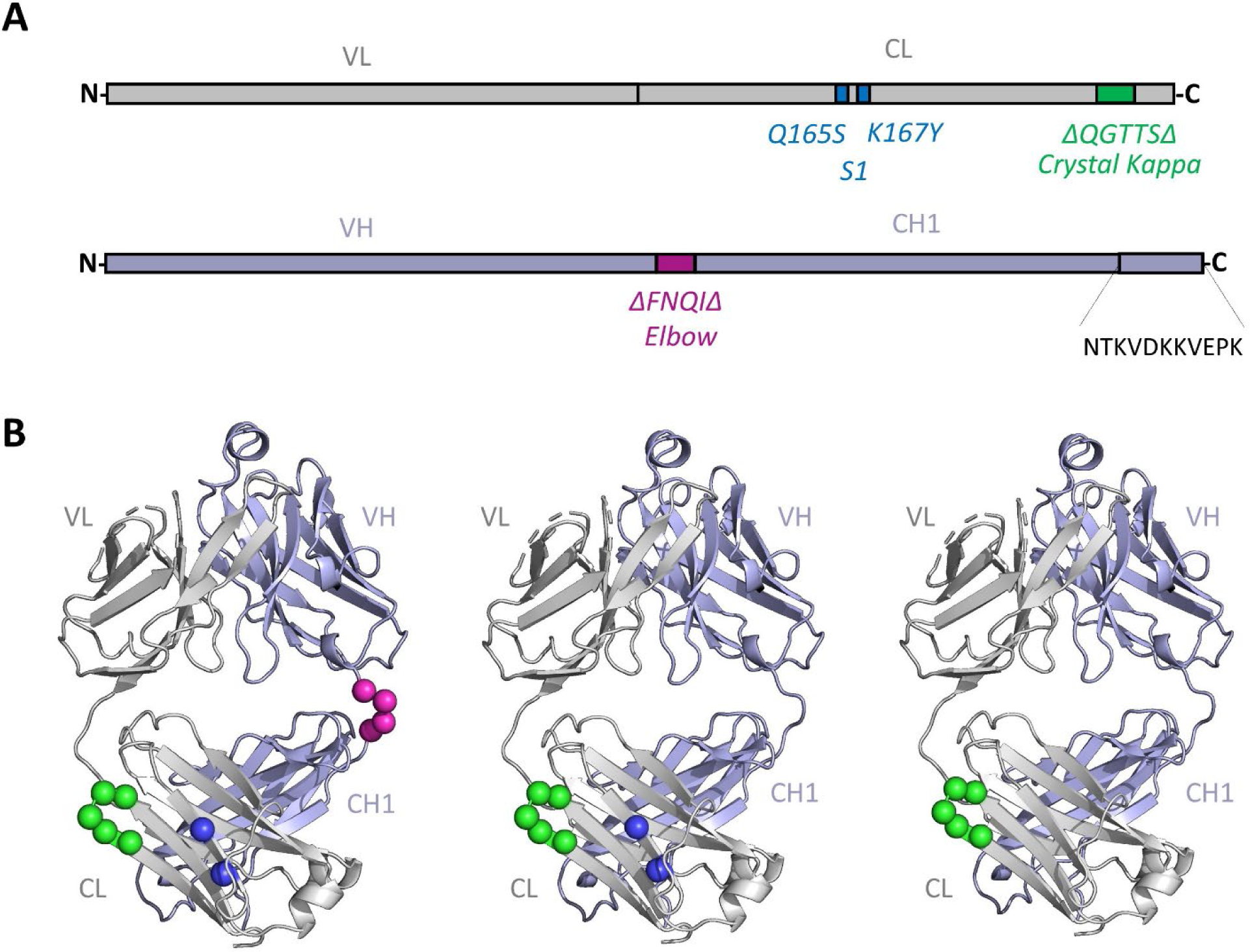
Fab frameworks with improved crystallisability. **(A)** Schematic representation of the Fab light- chain and heavy-chain (coloured grey or light blue, respectively) with the locations of S1 (blue), Crystal Kappa (green), and heavy-chain elbow (magenta) substitutions highlighted. To incorporate Crystal Kappa, residues HQGLSSP are substituted with QGTTS in the CL domain, whilst the C-terminus of the CH domain should include NTKVDKKVEPK, as described [21]. To incorporate the elbow substitution, SSAST is substituted with FNQI at the linker region connecting the VH and CH domains, as described [19]. **(B)** The three best Fab frameworks for enhancing crystallisation. The Fab^S1CE^ framework (left), which contains all three substitutions, should be prioritized, but given an adequate supply of antigen, Fab^S1C^ (centre) and Fab^C^ (right) should also be tried. Substituted regions are colored as in (**A**) and substituted residues are shown as spheres.

It is important to note that the S1 and Crystal Kappa substitutions reside in the constant region of the light chain, and thus, this mutation combination can be applied to any Fab of interest, regardless of source or species. By simply combining the VH and VL domains from a Fab of interest with the CL and CH domains reported here, chimeric Fabs with enhanced crystallisability can be created in a modular manner. Furthermore, whilst the elbow substitution has only been used thus far in the context of Fabs containing kappa light chains, it has potential to be used for crystallisation of Fabs containing lambda light chains as well. In sum, we anticipate that these strategies will greatly accelerate the resolution of Fab:antigen complexes of basic and therapeutic interest, and moreover, will further advance the use of Fabs as chaperones for the structural analysis of proteins and complexes that are recalcitrant to crystallisation.

## MATERIALS AND METHODS

### Construction and screening of phage-displayed libraries

Phage-displayed Fab libraries were constructed using a phagemid system, as described [25], with diversified positions and degenerate codons described in Figure 1. Phage pools representing the libraries were cycled through rounds of binding selections with EPHA2-FN2-Fc fusion protein immobilised on Maxisorp Immuno plates (ThermoFisher, #12-565-135), as described previously [21]. After five rounds of selection, individual clones were assayed for specific binding to EPHA2-FN2-Fc fusion protein, and positive clones were subjected to DNA sequence analysis.

### Fab protein production

For bacterial expression to screen Fab F1 SER framework variants, the genes encoding the Fab light and heavy chains were cloned into a bicistronic expression vector with F1 *ori*, Amp^R^ and lacIq components. *E.coli* BL21 (DE3) cultures harbouring the expression vector were grown to 0.6 OD_600_ in 2xYT media supplemented with 50 µg/mL of carbenicillin, followed by induction upon addition of 1 mM IPTG and incubation overnight at 18 °C. The cell pellet was harvested by centrifugation, resuspended, and lysed in lysis buffer (Phosphate Buffered Saline (PBS), 1% Triton X-100, 250 U/L benzonase, 2 mM MgCl_2_, 0.1 mM phenylmethylsulfonyl fluoride, and 1 g/L lysozyme). Cellular debris was removed by centrifugation. Fab protein was purified by rProtein A Sepharose (GE Healthcare) chromatography, and after elution with Pierce™ IgG Elution Buffer, was buffer exchanged into 20 mM HEPES pH 7.5, 100 mM NaCl followed by clarification by centrifugation.

For bacterial expression of Fabs, genes encoding the heavy and light chains were cloned into the pRH2.2 bicistronic expression vector suitable for bacterial expression and transformed into *E. coli* BL21 (DE3) competent cells. Cells were grown in 2xYT media containing 100 µg/mL of ampicillin at 37 °C for 2- 3 hours. Expression of Fabs was induced with 1 mM IPTG at 0.8-1.2 OD_600_, and cells were harvested by centrifugation after further growth for 3-4 hours after induction. Cells were homogenized in PBS supplemented with 1 mM PMSF. The cell lysate was incubated at 63 °C for 30 minutes prior to centrifugation that cleared lysate solution from cell debris. The cleared lysate was loaded onto a 5-mL HiTrap Protein-L affinity column. The column was pre-equilibrated and washed with buffer containing 20 mM Tris HCl pH 7.5 and 500 mM NaCl. Eluted Fabs in acetic acid from Protein-L column were loaded onto a 1-mL Resource S cation exchange column that was pre-equilibrated and later washed with buffer containing 50 mM Sodium Acetate pH 5.0. Fabs were eluted using a gradient of buffer containing 50 mM Sodium Acetate pH 5.0 and 2 M NaCl. Eluted fractions were concentrated against PBS.

For mammalian expression of Fabs, the genes encoding the heavy- and light-chains were cloned into separate vectors, suitable for mammalian expression [4]. Briefly, Expi293™ cell (ThermoFisher) culture was grown to a density of 2.6 x 10^6^ cell/ml in Expi293 media (Gibco) before co-transfection with Fab heavy- and light-chain expression vectors using FectoPRO® DNA transfection kit (Polyplus-transfection). Cells were kept at 37 °C, 8% CO_2_, 80% humidity with shaking at 125 rpm for 5-6 days to allow protein expression to proceed. Cells were pelleted by centrifugation and Fab protein was purified as described above for bacterial expression.

### Differential Scanning Fluorimetry

Melting temperature (°C) was determined by adding SYPRO™ Orange protein stain (Thermo Fisher) to 5 μM Fab protein in PBS and performing a thermal melt of 25-95 °C (0.5 °C /30 sec intervals), as described [27].

### Antigen protein preparation

The VHH domain was expressed and purified as described [24]. For other antigens, the gene encoding each antigen was cloned into a mammalian expression vector by bacterial homologous recombination [28]. To facilitate purification, a thrombin cleavage site followed by a hexa-histidine tag was fused to the C-terminus of EPHA2-FN2 or antigen-A, whereas a papain cleavage site followed by an Fc-tag was fused to the C-terminus of antigen B. The expression vector was transfected in mammalian cell culture using the Expi293 expression system (ThermoFisher # A14635), as described above. Cell culture expressing antigen-B was supplemented with 5 mM Kifunensine (MedChemExpress) to inhibit mannosidase I activity. EPHA2- FN2 and antigen-A were purified using His 60 Ni Superflow resin (Takara), whilst antigen-B was purified with rProteinA Sepharose (GE Healthcare). Affinity tags were cleaved by incubation with either thrombin (Merck) or papain (Thermo Scientific). Antigen B was further purified from Fc using rProteinA Sepharose (GE Healthcare), followed by deglycosylation with endoH (New England Biolabs). Purified protein was buffer exchanged into 20 mM HEPES pH 7.5, 100 mM NaCl, and clarified by centrifugation.

### Preparation of Fab:Antigen complexes for crystallisation

Protein purity and homogeneity was assessed using both denaturing and native polyacrylamide gel electrophoresis (Mini-PROTEAN TGX Stain-Free Precast Gels Bio-Rad). Fab^WT^ F1 and SER variants (S1-S7) were checked by size exclusion chromatography using a Superdex 200 Increase 10/300 column (Fisher Scientific) and monitoring elution at 215 nm. Fab-F1:EPHA2-FN2, Fab-14836:antigen-A, and Fab- 14836:antigen-B complexes were prepared for crystallisation screening at a 1:2 molar ratio, 7 mg/ml protein in 20 mM HEPES pH 7.5, 100 mM NaCl.

Fab-V1:VHH complexes were prepared at a 1:1.5 molar ratio and incubated at 4 °C for 2 hours, followed by purification using a Superdex 200 Increase 10/300 GL column. The eluted fractions containing Fab-V1:VHH complex in buffer (10 mM Tris-HCl pH 7.5, 150 mM NaCl) were well separated from excess of VHH domain, before pooling and concentrating to 7 mg/ml for crystallisation screening.

### Screening and optimisation of apo Fab and Fab:antigen complex crystallisation

Fab-V1:VHH complexes were crystallised by sitting drop vapor diffusion technique using Mosquito Crystal robot (SPT Labtech) at room temperature. Crystallisation was set up by mixing 0.1 µl of protein complex solution with 0.1 µl of screen solution from 50 µl reservoir solution on 96-well plate (TTP Labtech). In addition to JCSG-plus HT-96 Eco and PACT Premiere HT-96 (Molecular Dimensions), crystallization screens INDEX HT, PEG/Ion HT (Hampton Research) and ProComplex (NeXtal) were used, as specified in Supplementary Figure S7. Crystallization plates were incubated at 19 °C and manually checked by microscopy. In total, 119 crystals of various Fab variants complexed with VHH domain were supplemented with appropriate cryoprotectant (condition dependent) before being flash-frozen in liquid nitrogen for data collection.

For apo Fab and all other Fab:antigen complexes, a Mosquito Crystal robot (SPT Labtech) was used to set up sitting drop crystallisation screens with protein:precipitant drops at 0.2 µl:0.2 µl with 40 µl reservoir solution on 96-well plates (Hampton Research), at room temperature. Commercial screens JCSG- plus HT-96 Eco and PACT Premiere HT-96 (Molecular Dimensions), SaltRX HT, INDEX HT, GRAS Screen^TM^ 1 and GRAS Screen^TM^ 2 (Hampton Research) were used. Crystallization plates were incubated at room temperature and manually checked by microscopy.

Fab^S1^ F1 was subjected to refinement in Cryschem sitting drop 24-well plates (Hampton Research), for which plate-like crystals emerged using a crystallization liquor of 20% PEG 3350, 200 mM Ammonium Sulfate and 100 mM Bis Tris HCl pH 6.5, and from which a dataset at 3.5 Å was obtained. Fab^C^ F1 crystals were obtained using the sitting drop method with 22% PEG 8000, 200 mM NaCl and 100 mM sodium acetate pH 4.0 precipitant, and from which a 1.95 Å dataset was collected after cryoprotecting with 16% PEG8000, 100 mM sodium acetate pH 4.0, 200 mM NaCl. Fab^S1CE^ F1 crystals emerged from the broad crystallisation screening condition PACT, F5 (0.2 M sodium nitrate, 0.1 M Bis-Tris propane pH 6.5, 20 % PEG (w/v) 3350), from which one crystal diffracted to 2.6 Å after cryoprotecting with the precipitant supplemented with 25 % (v/v) ethylene glycol and flash freezing in liquid nitrogen. Fab^C^ -F1:EphA2-FN2 complex crystals were obtained using the sitting drop method in 24-well plates with 1.5 M sodium formate, 100 mM Tris HCl pH 8.0 precipitant. Prior to flash-freezing in liquid nitrogen, crystals were supplemented with 25% ethylene glycol for cryoprotection.

### Data collection and structure determination and refinement

For Fab-V1:VHH complexes, X-ray diffraction experiments were performed at beamline 24-ID-C or 24-ID- E at the Northeastern Collaborative Access Team (NECAT) at the Advance Photon Source at Argonne National Laboratory (Argonne IL). Datasets were collected remotely from a single crystal at 100 K using web-based remote GUI developed by NECAT team. Individual datasets were indexed, integrated with XDS [29] and scaled with Aimless [30]. Structures were solved by molecular replacement method using PHASER [31] with starting model of Fab^WT^-V1:VHH complex monomer (PDB ID: 7RTH)[24]. Structures were refined in PHENIX [32] or Refmac [33] and manually corrected in Coot [34]. The crystal contact and the surface of accessible solvent area analyses were performed by PISA [35]. Structural figures were made with CCP4mg [36].

For all other crystals, data collection was performed at Argonne National Laboratory at beamline 24-ID-E (NE-CAT). Datasets were collected remotely from a single crystal at 100 K using web-based remote GUI developed by NECAT team. Individual datasets were indexed, integrated with XDS [29] or Mosflm (T.G.G. Battye, L. Kontogiannis, O. Johnson, H.R. Powell and A.G.W. Leslie (2011), Acta Cryst. D67, 271- 281.), and scaled with Aimless [30]. The crystal structures of the Fab^S1^ F1, Fab^C^ F1, Fab^S1CE^ F1, and Fab^C^- F1:EPHA2-FN2 were solved from 3.5 Å, 1.9 Å, 2.6 Å and 4.2 Å datasets, respectively, by molecular replacement using PHASER [31]. Model refinement was undertaken using an iterative combination of manual re-modelling with Coot [34] and automated fitting and geometry optimization with Phenix.refine [32], including use of TLS parameters [37].

### Data deposition

Coordinates and structure factors have been deposited into the Protein Data Bank under the following accession codes: Fab^S1^ F1 (PDB: 8T7F), Fab^C^ F1 (PDB: 8T7G), Fab^S1CE^ F1 (PDB: 8T7I), Fab^C^-F1:EPHA2-FN2 (PDB: 8T9B), Fab^C^-V1:VHH (PDB: 8T6I), Fab^E^-V1:VHH (PDB: 8T58), Fab^CE^ -F1:VHH (PDB: 8T9Y), and Fab^S1CE^- F1:VHH (PDB: 8T8I).

## ACKNOWLEDGEMENTS

We are grateful to Lia Cardarelli, Greg Martyn and Shane Miersch for their valuable insights and helpful advice. This research was funded by Bristol-Myers-Squibb.

---

**Crystallography Data Table.**

|  | Fab<sup>S1</sup> F1 | Fab<sup>C</sup> F1 | Fab<sup>S1CE</sup> F1 | Fab<sup>C</sup> -F1:EPHA2-FN2 | Fab<sup>E</sup> -V1:VHH | Fab<sup>C</sup> -V1:VHH | Fab<sup>CE</sup> -V1:VHH | Fab<sup>S1CE</sup> -V1 :VHH |
| --- | --- | --- | --- | --- | --- | --- | --- | --- |
| Crystallization condition | 0.2 M Ammonium sulfate | 0.2 M NaCl | 0.2 M Sodium Nitrate | 1.5 M Sodium formate | 4 % Tacsimate pH 4.0 | 0.2 M Potassium citrate tribasic monohydrate |  | 2% Tacsimate pH 5.0 |
|  | 0.1 M Bis Tris HCl pH 6.5 | 0.1 M Sodium acetate pH 4.0 | 0.1 M Bis-Tris propane pH 6.5 | 0.1 M Tris HCl pH 8.0 | 12% PEG3350 |  | 0.1 M Citric acid pH 3.5 | 0.1 M Na citrate tribasic dihydrate pH 5.6 |
|  | 20 % PEG 3350 | 22 % PEG 8000 | 20 % PEG 3350 |  |  | 20% PEG 3350 | 25% PEG 3350 | 16% PEG 3350 |
| Freezing conditions | 25 % polyethyleneglycol | 25 % polyethyleneglycol | 25 % polyethyleneglycol | 25 % polyethyleneglycol | 10% Glycerol&25% PEG 1500 | 10% Glycerol | Crystallization solution | 30% Jeffamine |
| Space group | *P*2<sub>1</sub> 2<sub>1</sub> 2<sub>1</sub> | *C*2 | *P*4<sub>2</sub> 2<sub>1</sub> 2 | *P*2<sub>1</sub> | *P*2<sub>1</sub> | *P*2<sub>1</sub> 2<sub>1</sub> 2<sub>1</sub> | *P*4<sub>3</sub> 2<sub>1</sub> 2 | *P*2<sub>1</sub> 2<sub>1</sub> 2<sub>1</sub> |
| Unit cell parameters: |  |  |  |  |  |  |  |  |
| Length (Å) | a=80.8,<br>b=189.5,<br>c=216.5 | a=106.7,<br>b=73.8,<br>c=70.0 | a=74.0,<br>b=74.0,<br>c=219.1 | a=76.7,<br>b=130.7,<br>c=223.0 | a=65.8, b=119.8,<br>c=82.2 | a=54.5, b=74.2,<br>c=218.4 | a=b=73.6,<br>c=207.2 | a= 52.2, b=72.8,<br>c=242.4 |
| Angles (°) | α=β=γ=90.0 | α=γ=90.0<br>β=116.5 | α=β=γ=90.0 | α=γ=90.0<br>β=89.8 | α=γ=90, β=93.35 | α=β=γ=90 | α=β=γ=90 | α=β=γ=90 |
| Matthews coefficient (Å<sup>3</sup>/Da) | 2.84 | 2.53 | 3.08 | 4.76 | 2.5 | 3.5 | 2.3 | 3.7 |
| Solvent content (%) | 56.6 | 51.4 | 60 | 74.2 | 50.7 | 64.9 | 46.7 | 66.9 |
| Data collection: |  |  |  |  |  |  |  |  |
| Resolution (Å) | 95.04-3.50<br>(3.64-3.50) | 62.65-1.84<br>(1.88-1.84) | 74.0-2.50<br>(2.60-2.50) | 131.33-4.20<br>(4.43-4.20) | 119.83-2.23<br>(2.29-2.23) | 74.19-2.55(2.66-2.55) | 103.62-2.52<br>(2.62-2.52) | 121.2-2.52<br>(2.62-2.52) |
| Number of unique reflections | 40200 (4729) | 40653 (1987) | 20950 (2408) | 31847 (4648) | 61587 (4313) | 29702 (3539) | 20161 (2220) | 32276 (3603) |
| Completeness (%) | 93.1 (99.0) | 96.4 (78.1) | 95.0 (100) | 97.3 (97.8) | 99.1 (93.8) | 99.7 (99.9) | 100 (99.7) | 100 (100) |
| Avg (I/σ(I)) | 7.9 (2.1) | 12.7 (1.1) | 8.1 (1.5) | 4.2 (1.2) | 16.1 (2.7) | 18.5 (1.0) | 19.2 (1.1) | 12.5 (0.8) |
| R<sub>pim</sub> | 0.07(0.41) | 0.058(0.87) | 0.07(0.64) | 0.12(0.81) | 0.03 (0.24) | 0.03 (0.8) | 0.03 (0.51) | 0.04 (0.78) |
| Redundancy | 6.7(6.7) | 3.8(2.7) | 6.4(6.7) | 3.0(3.0) | 6.9 (6.2) | 5.6 (5.8) | 25.9 (26.1) | 13.1 (13.6) |
| CC<sub>1/2</sub> | 1.00(0.79) | 1.00(0.37) | 1.00(0.63) | 1.00(0.60) | 0.99 (0.86) | 0.99 (0.54) | 0.99 (0.69) | 0.99 (0.37) |
| Wilson B-factor (Å<sup>2</sup>) | 91.7 | 33.6 | 42.7 | 101.4 | 46.5 | 87.2 | 74.7 | 80.6 |
| **Refinement:** |  |  |  |  |  |  |  |  |
| Fab/Fab:antigen complex molecules in the ASU | 6 | 1 | 1 | 4 | 2 | 1 | 1 | 1 |
| R<sub>work</sub> / R<sub>free</sub> | 0.242/0.306 | 0.194/0.228 | 0.200/0.256 | 0.296/0.315 | 0.18/0.225 | 0.194/0.254 | 0.214/0.285 | 0.191/0.243 |
| **No. atoms** |  |  |  |  |  |  |  |  |
| Protein/peptide | 18925 | 3157 | 3147 | 15657 | 8611 | 4267 | 4121 | 4319 |
| Ligand/ion | 40 | 21 | 15 | — | 14 | 6 | 14 | 27 |
| Water | — | 214 | 83 | — | 569 | 39 | 37 | 71 |
| **B-factors** |  |  |  |  |  |  |  |  |
| Overall | 96.6 | 46.2 | 62.6 | 156.3 | 50.1 | 97.2 | 86.5 | 83.2 |
| Protein | 96.6 | 46.2 | 63.2 | 156.3 | 49.9 | 97.1 | 88.9 | 83.3 |
| Ligand/ion | 72.7 | 51.7 | 52.4 | — | 57.8 | 110.1 | 107.9 | 105 |
| Water | — | 45.4 | 43.4 | — | 49.8 | 98.7 | 69.6 | 75 |
| **R.m.s. deviations** |  |  |  |  |  |  |  |  |
| Bond lengths (Å) | 0.005 | 0.007 | 0.008 | 0.005 | 0.004 | 0.008 | 0.004 | 0.004 |
| Bond angles (°) | 1.00 | 0.86 | 1.07 | 1.01 | 1.04 | 1.10 | 1.05 | 1.12 |
| **Ramachandran** |  |  |  |  |  |  |  |  |
| Favoured | 91.3 | 97.3 | 96.2 | 85.9 | 96.9 | 95.1 | 92.9 | 94.5 |
| Allowed | 8.4 | 2.7 | 3.8 | 13.2 | 3.1 | 4.72 | 6.91 | 5.5 |
| Forbidden | 0.3 | 0 | 0 | 0.9 | 0 | 0.18 | 0.19 | 0 |

