## Supplementary_Material for "Engineered Antigen-Binding Fragments for Enhanced Crystallisation of Antibody:Antigen Complexes"

| **Number** | **PDB ID** | **Resolution (Å)** | **Antigen bound or in Fab apo form** |
| --- | --- | --- | --- |
| 1 | 1B2W | 2.90 | - |
| 2 | 1BJ1 | 2.40 | Vascular endothelial growth factor (VEGF) |
| 3 | 1JPS | 1.85 | Tissue factor |
| 4 | 1N8Z | 2.52 | HER2 |
| 5 | 1S78 | 3.25 | ErbB2 |
| 5 | 1TZH | 2.60 | h-VEGF |
| 6 | 1TZI | 2.80 | h-VEGF |
| 7 | 1ZA3 | 3.35 | Death receptor 5 (DR5) |
| 8 | 2FGW | 3.00 | - |
| 9 | 2FJH | 3.10 | VEGF |
| 10 | 2H9G | 3.23 | DR5 |
| 11 | 2HFF | 1.95 | - |
| 12 | 2HFG | 2.61 | hBR3 |
| 13 | 2JIX | 3.20 | Soluble domain of EPO receptor |
| 14 | 2QQL | 3.10 | Neuropilin-2 a1a2b1b2 Domains |
| 15 | 2QQN | 2.20 | Neuropilin-1 b1 Domain |
| 16 | 2QR0 | 3.50 | VEGF |
| 17 | 2R8S | 1.95 | P4-P6 RNA ribozyme domain |
| 18 | 2WUB | 2.90 | Hepatocyte growth factor activator (HGFA) |
| 19 | 2WUC | 2.70 | HGFA |
| 20 | 3BN9 | 2.17 | MT-SP1 |
| 21 | 3DVG | 2.60 | K63-linked di-ubiquitin |
| 22 | 3DVN | 2.70 | K63-linked di-ubiquitin |
| 23 | 3EFD | 2.60 | Cytoplasmic domain of KcsA |
| 24 | 3EFF | 3.80 | KcsA in its closed conformation |
| 25 | 3EO9 | 1.80 | - |
| 26 | 3EOA | 2.80 | LFA-1 I domain, Form I |
| 27 | 3EOB | 3.60 | LFA-1 I domain, Form I |
| 28 | 3HC3 | 1.72 | - |
| 29 | 3IDX | 2.50 | HIV-gp120 core |
| 30 | 3JWD | 2.61 | HIV-1 gp120 with gp41-Interactive Region |
| 31 | 3KR3 | 2.20 | IGF-II |
| 32 | 3L95 | 2.19 | human Notch1 Negative Regulatory Region (NRR) |
| 33 | 3N85 | 3.20 | HER2 extracellular regions |
| 34 | 3PGF | 2.10 | Maltose binding protein (MBP) |
| 35 | 3PJS | 3.80 | Full-Length KcsA K+ Channel |
| 36 | 3PNW | 2.05 | Tudor domain of human TDRD3 |
| 37 | 3R1G | 2.80 | BACE1 |
| 38 | 3SKJ | 2.50 | Human ephrin type A receptor 2 (EphA2) |
| 39 | 3U30 | 2.43 | Linear Ubiquitin |
| 40 | 3BDY | 2.60 | VEGF |
| 41 | 3SOB | 1.90 | Lrp6 |
| 42 | 3K2U | 2.35 | HGFA |
| 43 | 2R56 | 2.80 | Bovine Beta-Lactoglobulin Allergen |

**Supplementary Table S1. Fab and Fab:antigen complex crystal structures.** The listed PDB entries were processed by the CryCo server [1] to evaluate Fab surface residue participation in crystal lattice contact interfaces.

| **Resolution**  **Space group**  **a b c**  **α β γ** | **Fab^S1^ F1 (3.5 Å)**  ***P*2_1_2_1_2_1_**  **80.7, 189.4, 216.5**  **90, 90, 90** | **Fab^C^ F1 (1.9 Å)**  ***C*2**  **106.7, 73.8, 70.0**  **90.0, 116.5, 90.0** | **Fab^S1CE^ F1 (2.6 Å)**  ***P*4_2_2_1_2**  **74.0, 74.0, 219.1**  **90.0, 90.0, 90.0** |
| --- | --- | --- | --- |
| **Chain A**  **(Heavy chain)** | 74A : 206H (-x, y+1/2, -z+1/2)  117A : 117K (x, y, z) | 1, 26A : 180A (-x, y, -z)  16, 93A : 7, 8B (-x+1/2, y+1/2, -z)  27A : 170B (x+1/2, y+1/2, z)  29A : 165, 167B (x+1/2, y+1/2, z)  60, 62A : 60, 62A (-x, y, -z)  85A : 172B (x+1/2, y+1/2, z)  140A : 157A (-x+1/2, y+1/2, -z)  157A : 140A (-x+1/2, y+1/2, -z)  180A : 1A (-x, y, -z)  182A : 70B (-x, y, -z)  214A : 28B (x, y, z)  228-236A : 218-226B (-x+1/2, y+1/2, -z) | 14, 136-139A : 152-156, 208A (-x+1/2, y+1/2, -z+1/2)  59, 60, 82, 111, 112A : 59, 60, 82, 111, 112A (-y, -x, z)  152-156, 182A : 14, 136-139A (-x+1/2, y+1/2, -z+1/2)  179-181A : 179-181A (-x, -y, z)  184A : 170B (x+1/2, -y+1/2, -z+1/2)  195A : 195A (-x, -y, z)  227-236A : 218-227B (-x+1/2, y+1/2, -z+1/2) |
| **Chain B**  **(Light chain)** | 167, 169-171B : 15, 16, 93, 95A (x+1/2, -y+1/2, -z) | 7, 8B : 16, 93A (-x+1/2, y+1/2, -z)  28B : 214A (x, y, z)  66B : 80B (-x, y, -z)  69, 70B : 179, 182A (-x, y, -z)  165, 167B : 29A (x+1/2, y+1/2, z)  170-172B : 27-30A (x+1/2, y+1/2, z)  218-227B : 228-236A (-x+1/2, y+1/2, -z) | 144-146B : 200-203, 206B (-x, -y, z)  167, 170, 172, 174B : 156, 184, 208, 209, 210A (x+1/2, -y+1/2, -z+1/2)  200-203B : 145, 146B (-x, -y, z)  218-227B : 228-234A (-x+1/2, y+1/2, -z+1/2) |
| **Chain C (HC)** | 116, 117, 120C : 29, 116, 117, 120F (x, y, z) | **-** | **-** |
| **Chain D (HC)** | 15, 16D : 170, 171G (x+1/2, -y+1/2, -z)  93, 95D : 169, 170G  116, 117D : 117, 120I (x, y, z)  120D : 29J (x, y, z)  214D : 162L (-x, y+1/2, -z+1/2)  179, 180, 219D : 167, 172L (-x, y+1/2, -z+1/2) | **-** | **-** |
| **Chain E (LC)** | 28-30E : 28, 80, 84J (x, y, z)  84, 85E : 83, 84J (x, y, z)  174E : 215K (-x, y+1/2, -z+1/2) | **-** | **-** |
| **Chain F (HC)** | 117F : 112, 116, 117C (x, y, z)  116F : 116I (x, y, z)  116F : 112, 117K (x, y, z)  215F : 174J (-x, y+1/2, -z+1/2) | **-** | **-** |
| **Chain G (LC)** | 28-30G : 28, 31, 80, 84H (x, y, z)  57G : 28H (x, y, z)  84, 85G : 83H (x, y, z)  169-171G : 15, 93, 95D (x+1/2, -y+1/2, -z) | **-** | **-** |
| **Chain H (LC)** | 29H : 117, 120C (x, y, z)  28H : 57G (x, y, z)  29H : 115K (x, y, z)  83, 86H : 84, 85G (x, y, z)  84, 85H : 30, 89G (x, y, z)  169, 170H : 16A (-x, y+1/2, -z+1/2)  172, 173H : 219, 180I (-x, y+1/2, -z+1/2)  206, 207H : 74, 92A (-x, y+1/2, -z+1/2) | **-** | **-** |
| **Chain I (HC)** | 15, 16, 93, 95I : 167, 169-172B (x+1/2, -y+1/2, -z)  116I : 116F (x, y, z)  116I : 112A (x, y, z)  117I : 112, 116, 117D (x, y, z)  180I : 172H (-x, y+1/2, -z+1/2) | **-** | **-** |
| **Chain J (LC)** | 29J : 117, 120D (x, y, z)  29J : 115A (x, y, z)  28J : 30E (x, y, z)  80J : 28, 29E (x, y, z)  165, 167, 172, 173, 174J : 215, 216F (-x, y+1/2, -z+1/2) | **-** | **-** |
| **Chain K (HC)** | 115K : 29H (x, y, z)  117K : 117A (x, y, z)  117K : 115F (x, y, z) | **-** | **-** |
| **Chain L (LC)** | 80L : 84B (x, y, z)  144L : 5, 24E (-x y+1/2, -z+1/2)  161, 162, 163, 165, 167, 172, 174 L : 214, 219, 180, 182D (-x, y+1/2, -z+1/2) | **-** | **-** |

**Supplementary Table S2. Crystal lattice contact interactions for Fab^S1^ F1, Fab^C^ F1 and Fab^S1CE^ F1 ASUs.** NB: packing interactions defined manually in *C*oot [2], with IMGT numbering used [3]. PDB chain IDs are shown in the first column. Highlighted in yellow are the S1 substitutions Q165S/K167Y and other neighbouring residues (165-174) in the light chain CH domain involved in the same crystal lattice packing interaction site. Highlighted in grey, are the non-mutated Q165/K167 residues and other light chain CH domain residues (165-174) involved in the same crystal lattice packing interaction. Highlighted in cyan are residues involved in the Crystal Kappa-mediated packing interaction. Highlighted in red are elbow (WT or substituted) residues involved in crystal lattice packing in the elbow junction, proximal to the Crystal Kappa packing site.

| **Resolution**  **Space group**  **a b c**  **α β γ** | **Fab^C^-F1:EPHA2-FN2 (4.2 Å)**  ***P*2_1_**  **76.7, 130.7, 223.0**  **90, 89.8, 90** |
| --- | --- |
| **Chain A (Heavy chain)** | 82A : 465L (x, y, z)  156A : 227D (-x, y+1/2, -z)  228-234A : 218-226E (-x, y+1/2, -z) |
| **Chain B (Light chain)** | 218-226B : 228-234D (-x, y+1/2, -z) |
| **Chain C (HC)** | 81-83C : 460, 463K (x, y, z)  228-234C : 218-226H (-x, y+1/2, -z) |
| **Chain D (HC)** | 228-234D : 218-226B (-x, y+1/2, -z) |
| **Chain E (LC)** | 218-226E : 226-234A (-x, y+1/2, -z) |
| **Chain F (HC)** | 82, 83F : 460, 463I (x, y, z)  228-234F : 218-226G (-x, y+1/2, -z) |
| **Chain G (LC)** | 218-226G : 228-234F (-x, y+1/2, -z) |
| **Chain H (LC)** | 218-226H : 228-234C (-x, y+1/2, -z) |

**Supplementary Table S3. Crystal lattice contact interactions for Fab^C^-F1:EPHA2-FN2 ASU.** NB: packing interactions defined manually in *C*oot [2], with IMGT numbering used [3]. PDB chain IDs are shown in the first column. Highlighted in cyan are residues involved in the Crystal Kappa-mediated packing interaction. All crystal lattice packing interaction mediated by Fab Light or heavy chains in the ASU, with neighbouring molecules, are shown. Crystal lattice packing interaction mediated by the antigen EPHA2-FN2 in the ASU, with neighbouring molecules, are not included.

| **Resolution**  **Space group**  **a b c**  **α β γ** | **Fab^E^ V1:VHH (2.23 Å)**  ***P*2_1_**  **65.8, 119.8, 82.2**  **90.0, 93.4, 90.0** | **Fab^C^ V1:VHH (2.55 Å)**  ***P*2_1_2_1_2_1_**  **54.5, 74.2, 218.4**  **90.0, 90.0, 90.0** | **Fab^CE^ V1:VHH (2.52 Å)**  ***P*4_3_2_1_2**  **73.6, 73.6, 207.2**  **90.0, 90.0, 90.0** | **Fab^S1CE^ V1:VHH (2.52 Å)**  ***P*2_1_2_1_2_1_**  **52.2, 72.8, 242.4**  **90.0, 90.0, 90.0** |
| --- | --- | --- | --- | --- |
| **Chain A**  **(Antigen/VHH domain for all)** | 14A : 14D (x, y, z)  A15A : 206B (x, y, z)  18A : 80E (x, y, z)  20A : 117F (x, y, z)  20A : 30E (x, y, z)  27, 28, 30, 31A : 209, 224, 225, 226, 228E (-x, y+1/2, -z)  62, 64, 67, 72, 113A : -2, -1F (x, y, z)  69, 72, 74A : 144, 145B (x, y, z)  90A : 30E (x, y, z)  90A : 117F (x, y, z)  92A : 84, 85E (x, y, z)  93A : 200, 202B (x, y, z)  95A : 201B (x, y, z)  108A : 231, 228E (-x, y+1/2, -z)  108A : 153F (-x, y+1/2, -z)  110A : 120D (x, y, z)  112, 114A : 49, 118D (x, y, z)  113A : -1F (x, y, z)  115A : 115D (x, y, z)  111, 120, 121, 125A : 114, 115D (x, y, z) | 12, 14, 136, 137A : 114, 115A (x+1/2, -y+1/2, -z)  57, 60, 63-65, 110, 113, 160, 162A : 29-30, 80, 84, 85L ( x+1/2, -y+1/2, -z)  69A : 116, 118H (x+1/2, -y+1/2, -z)  72A : 113, 115H (x+1/2, -y+1/2, -z)  83, 84A : 70, 72, 74L (x+1/2, -y+1/2, -z)  114, 115A : 12, 14, 136, 137A (x+1/2, -y+1/2, -z)  64, 117A : 117H (x+1/2, -y+1/2, -z)  132A : 69A (x+1/2, -y+1/2, -z) | 14, 117, 79, 80, 83A : 137, 161, 162, 165, 167, 172, 174, 212, 213L (y, x, -z)  15, 16A : 180H (-y+1/2, x+1/2, z+3/4)  31, 57, 69A : 69, 72, 75, 93, 95H (-y+1/2, x+1/2, z+3/4)  58, 59, 62-74A : 60-67, 72A (-y, -x, -z+1/2)  95A : 182, 216H (-y+1/2, x+1/2, z+3/4)  114A : 24L (-x+1/2, y+1/2, -z+3/4)  125A : 95H (-y+1/2, x+1/2, z+3/4)  136, 137A : 95, 163, 165L (y, x, -z) | 12, 14, 137A : 114, 115A (x+1/2, -y+1/2, -z)  57, 60-64, 113A : 28, 30, 80, 84, 85L ( x+1/2, -y+1/2, -z)  64, 67, 69, 72A : 113, 117, 118, 132H (x+1/2, -y+1/2, -z)  81-83A : 70, 72L (x+1/2, -y+1/2, -z)  113A : 80, 84L (x+1/2, -y+1/2, -z)  114, 115A : 12, 14, 136, 137A (x+1/2, -y+1/2, -z)  117A : 117H (x+1/2, -y+1/2, -z)  9, 118, 131, 132A : 69A (x+1/2, -y+1/2, -z)  14, 136, 137A : 114, 115A (x+1/2, -y+1/2, -z) |
| **Chain B (LC)** | 144, 145B : 69, 72A (x, y, z)  200, 201, 202, 206B : 15, 93, 95A (x, y, z) | **-** | **-** | **-** |
| **Chain C (HC)** | 117C : 13, 17, 18D (x, y, z)  225, 228C : 3, 5F (x, y, z) | **-** | **-** | **-** |
| **Chain D (Antigen)** | 14D : 12, 14A (x, y, z)  17, 18D : 117C (x, y, z)  114D : 109, 120, 122, 125A (x, y, z)  115, 118, 120D : 112, 114, 115A (x, y, z) | **-** | **-** | **-** |
| **Chain E (HC)** | 30, 117^E^ : 20, 90A (x, y, z)  80, 84, 85^E^ : 18, 92A (x, y, z)  209, 226, 228, 231^E^ : 27, 28, 108A (-x, y+1/2, -z) | **-** | **-** | **-** |
| **Chain F (LC)** | -2, -1F : 62-64, 67, 72, 113A (x, y, z)  3, 5F : 225, 228C (x, y, z)  83F : 209, 211F (-x, y+1/2, -z)  129F : 230C (x, y, z)  153, 155, 156F : 33, 57, 59, 110A (-x, y+1/2, -z)  209, 211F : 83F (-x, y+1/2, -z)  227, 228F : 3, 5C (x, y, z) | **-** | **-** | **-** |
| **Chain L (LC)** | **-** | 27-30, 185L : 57, 60-64A (x+1/2, -y+1/2, -z)  70-72L : 83, 84A (x+1/2, -y+1/2, -z)  84, 85L : 62, 64, 110, 113A (x+1/2, -y+1/2, -z)  127, 128L : 144L (x+1/2, -y+1/2, -z)  144L : 127, 128L (-x, y+1/2, -z+1/2)  218-226L : 228-233H (-x, y+1/2, -z+1/2) | 18, 92L : 236H (-y+1/2, x+1/2, z+3/4)  72-75, 77L : 214-217, 234H (-y+1/2, x+1/2, z+3/4)  79, 80, 83, 84L : 162, 163, 179L (y, x, -z)  126L : 144L (x+1/2, -y+1/2, -z+1/4)  144L : 128L (x+1/2, -y+1/2, -z+1/4)  161-163, 165, 172, 174L : 12, 14, 80, 83, 117, 137L (y, x, -z)  218-227L : 228-236H (x+1/2, -y+1/2, -z+1/4) | 27, 28, 30, 85L : 57, 60, 61, 63, 64, 162A (x+1/2, -y+1/2, -z)  70, 72L : 81-83 (x+1/2, -y+1/2, -z)  84, 85L : 57, 60, 110, 113A (x+1/2, -y+1/2, -z)  127, 128L : 114L (-x, y+1/2, -z+1/2)  144L : 128L (-x, y+1/2, -z+1/2)  218-226L : 228-234, 236H (-x, y+1/2, -z+1/2) |
| **Chain H**  **(HC)** | **-** | 113, 114, 116, 118H : 69, 72A (x+1/2, -y+1/2, -z)  117H : 64, 114A (x+1/2, -y+1/2, -z)  227-234H : 218-226L (-x, y+1/2, -z+1/2) | 72H : 69A (-x+1/2, y+1/2, -z+3/4)  75, 93H : 31A (y+1/2, -x+1/2, z+1/4)  95H : 125A (y+1/2, -x+1/2, z+1/4)  117H : 174L (y, x, -z)  180, 182H : 15, 95A (y+1/2, -x+1/2, z+1/4)  215, 216H : 72, 74L (y+1/2, -x+1/2, z+1/4)  228-236H : 218-226L (x+1/2, -y+1/2, -z+1/4)  236H : 18L (y+1/2, -x+1/2, z+1/4) | 117, 118H : 64, 67, 69 (x+1/2, -y+1/2, -z)  137-139, 141, 227, 228H : 153-156H (-x, y+1/2, -z+1/2)  153-156H : 138, 139, 141, 226, 228H (-x, y+1/2, -z+1/2)  228-234, 236H : 218-226L (-x, y+1/2, -z+1/2) |

**Supplementary Table S4. Crystal lattice contact interactions for Fab^E^ V1:VHH, Fab^c^ V1:VHH, Fab^S1C^ V1:VHH, and Fab^S1CE^ V1:VHH ASUs.** PDB chain IDs are shown in the first column. IMGT numbering used [3]. Highlighted in grey are the non-mutated Q165/K167 residues and other residues involved in the same crystal lattice packing interaction. Highlighted in cyan are residues involved in the Crystal Kappa-mediated packing interaction. Highlighted in red are elbow (WT or substituted) residues involved in crystal lattice packing in the elbow junction, proximal to the Crystal Kappa packing site.

| **IMGT numbering** | **163** | **165** | **167** | **183** | **184** | **187** | **201** | **203** | **205** | **206** | **208** | **217** | **Yield**  **mgL^-1^** | **Tm**  **(°C)** | **Tm**  **std** |
| --- | --- | --- | --- | --- | --- | --- | --- | --- | --- | --- | --- | --- | --- | --- | --- |
| **WT sequence** | K | Q | K | E | Q | K | K | D | E | K | K | Q | 10.2 | 75.25 | 0.35 |
| **S1** | - | S | Y | - | - | - | - | - | - | - | - | A | 12.0 | 75.50 | 0.00 |
| **S2** | S | S | Y | - | - | - | - | - | - | - | - | A | 9.0 | 75.25 | 0.35 |
| **S3** | S | S | Y | - | - | - | S | - | Y | - | - | A | 7.9 | 75.25 | 0.35 |
| **S4** | S | S | Y | - | - | - | T | - | Y | - | - | A | 9.6 | 75.25 | 0.35 |
| **S5** | S | S | Y | - | - | - | T | - | - | S | - | A | 8.2 | 75.50 | 0.00 |
| **S6** | S | S | Y | - | - | - | T | - | Y | - | S | A | 10.4 | 75.25 | 0.35 |
| **S7** | S | S | Y | - | - | - | T | - | - | S | S | A | 8.3 | 75.50 | 0.00 |
| **S8** | S | S | Y | - | - | - | S | - | Y | S | - | A | 4.0 | 75.50 | 0.00 |
| **S9** | S | S | Y | - | - | - | T | - | Y | S | - | A | 7.8 | 75.25 | 0.35 |
| **S10** | S | S | Y | - | - | - | S | - | - | S | - | A | 6.3 | 75.25 | 0.35 |
| **S11** | S | S | Y | - | - | - | S | - | Y | - | S | A | 9.2 | 75.50 | 0.71 |
| **S12** | S | S | Y | - | - | - | S | - | - | - | S | A | 4.9 | 75.50 | 0.00 |
| **S13** | S | S | Y | - | - | - | T | - | - | - | S | A | 0.9 | 75.50 | 0.00 |
| **S14** | S | S | Y | - | - | - | S | - | Y | S | S | A | 4.0 | 75.50 | 0.00 |
| **S15** | S | S | Y | - | - | - | T | - | Y | S | S | A | 6.5 | 75.50 | 0.00 |
| **S16** | S | S | Y | - | - | - | S | - | - | S | S | A | 6.3 | 75.50 | 0.00 |
| **S17** | S | S | Y | - | - | - | A | - | A | A | A | A | 4.1 | 75.00 | 0.00 |
| **S18** | S | S | Y | Y | - | T | - | - | - | - | - | A | 13.9 | 73.25 | 0.35 |
| **S19** | S | S | Y | A | - | T | - | - | - | - | - | A | 8.8 | 73.50 | 0.71 |
| **S20** | S | S | Y | S | - | T | - | - | - | - | - | A | 13.3 | 72.75 | 0.35 |
| **S21** | S | S | Y | Y | P | T | - | - | - | - | - | A | 6.5 | 71.25 | 0.35 |
| **S22** | S | S | Y | A | P | T | - | - | - | - | - | A | 4.9 | 70.50 | 0.00 |
| **S23** | S | S | Y | S | P | T | - | - | - | - | - | A | 5.7 | 70.25 | 0.35 |
| **S24** | S | S | Y | Y | - | S | - | - | - | - | - | A | 6.9 | 72.75 | 0.35 |
| **S25** | S | S | Y | A | - | S | - | - | - | - | - | A | 5.4 | 71.25 | 0.35 |
| **S26** | S | S | Y | S | - | S | - | - | - | - | - | A | 13.3 | 72.75 | 0.35 |
| **S27** | S | S | Y | Y | P | S | - | - | - | - | - | A | 4.4 | 71.25 | 0.35 |
| **S28** | S | S | Y | A | P | S | - | - | - | - | - | A | 5.1 | 70.25 | 0.35 |
| **S29** | S | S | Y | S | P | S | - | - | - | - | - | A | 8.3 | 69.75 | 0.35 |

**Supplementary Figure S1.** **Fab F1 SER-variants screened for protein yield and thermostability.** The WT Fab sequence is shown at the top, numbered according to IMGT nomemclature [3], and the sequences of twenty-nine Fab F1 SER-variants (S1-29) are shown below. Dashes indicate identity with the WT sequence. The yield of protein from 1-L of bacterial culture and the T_m_ determined by differential scanning fluorimetry are shown to the right.

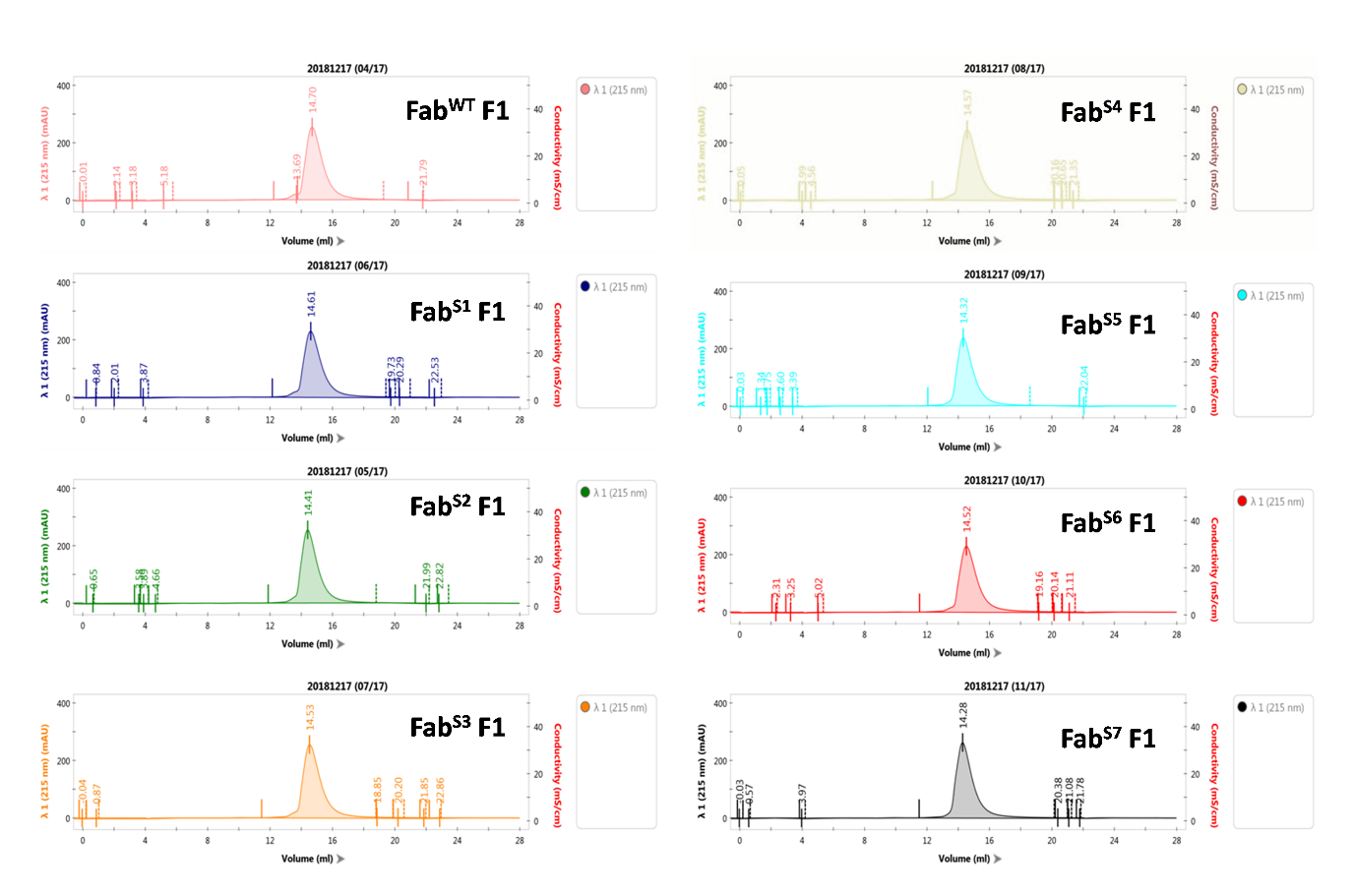

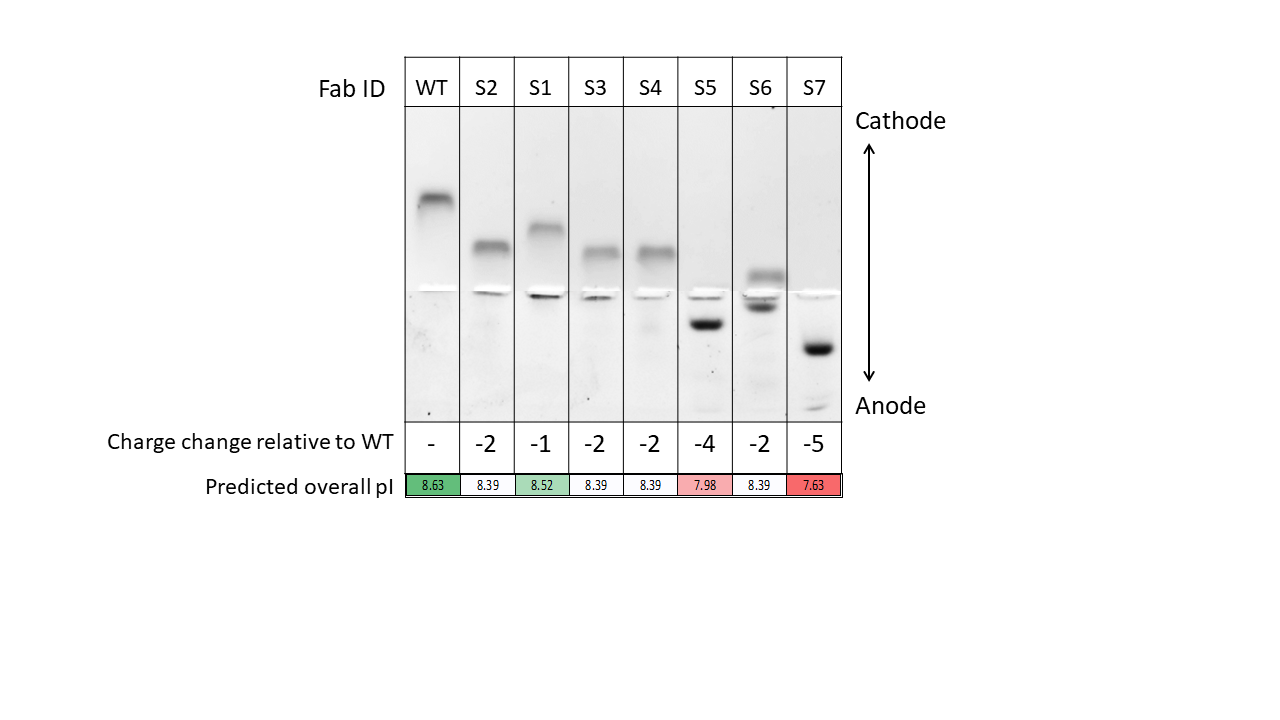

A

B

**Supplementary Figure S2. Purity and quality assessment of EPHA2-FN2 binding-Fab F1 and the seven most promising SER-modified Fab F1 variants (S1-S7).** **(A)** Size exclusion chromatography (SEC) elution profiles for each Fab monitored at a wavelength of 215 nm (x-axis = elution volume, y-axis = wavelength). Each Fab eluted as a single peak. **(B)** Polyacrylamide native gel electrophoresis was employed to assess the quality of the Fab proteins, prior to crystallisation screening. A single, discrete band can be observed for each Fab indicating compositional homogeneity. Furthermore, an internal comparison between the Fabs can be made based on their relative mobilities which are proportional to their respective predicted overall isoelectric point (pI) values. NB: two gels were performed; the first with the electric current directed to the cathode (shown above - flipped vertically), the second with electric current directed to the anode (below).

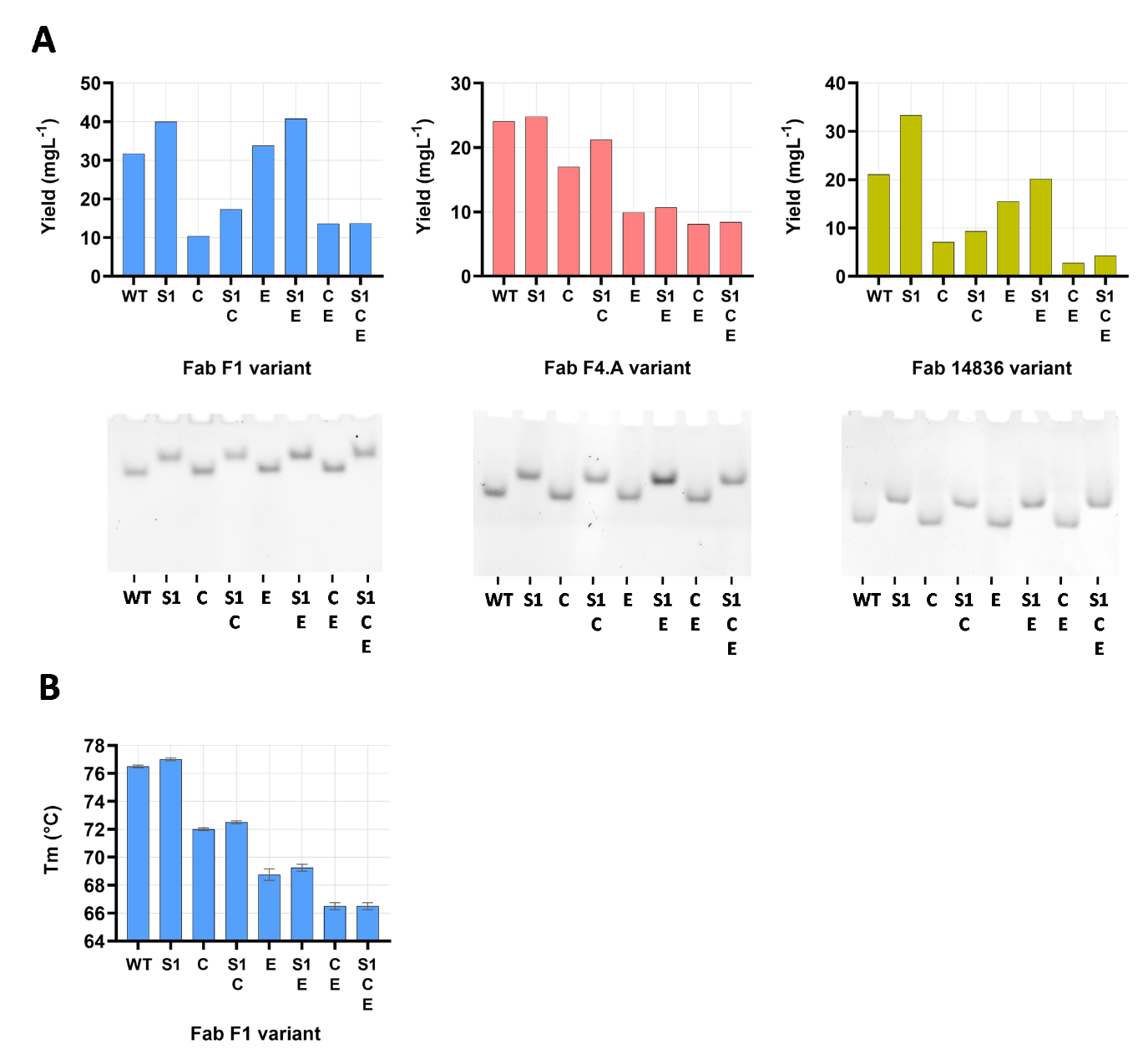

**Supplementary Figure S3. Yield, purity and quality assessment of in-house Fabs (IDs: F1, F4.A, 14836) with S1, Crystal Kappa and elbow mutations.** **(A)** Comparison of yield between WT Fab, and S1 +/- Crystal Kappa (C) +/- elbow (E) variants, is shown in top panel. In the lower panel, Native polyacrylamide gel electrophoresis was employed to determine the purity and compositional homogeneity of the Fab samples, prior to broad crystallisation screening. All Fabs run as a single, discrete band, indicating compositional homogeneity and monodispersity. NB: S1 substitutions reduces the pI of the Fab, resulting in a slower migration towards the cathode, resulting in a staggered Fab band migration pattern on the gels. **(B)** Differential scanning fluorimetry results for Fab^WT^ F1 and variants reveals that whilst the addition of Crystal Kappa and elbow substitutions reduces the Tm, the inclusion of S1 substitutions confers a marginal increase in Tm.

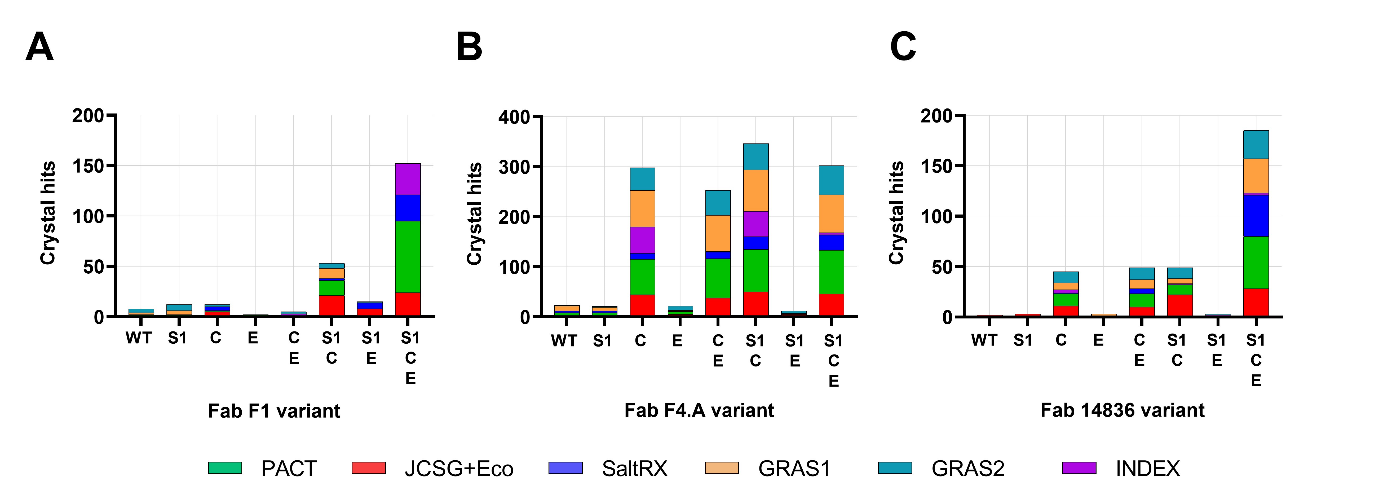

**Supplementary Figure S4. Results of crystallisation screens for apo Fab variants.** Results are shown for WT and indicated S1, Crystal Kappa (C), and elbow (E) variants (and their combinations) of **(A)** Fab F1, **(B)** Fab F4.A, and **(C)** Fab 14836. The number of crystal hits (y-axis) obtained for each Fab variant (x-axis) are shown after screening a total of 576 conditions (96 conditions for each of the six indicated screens).

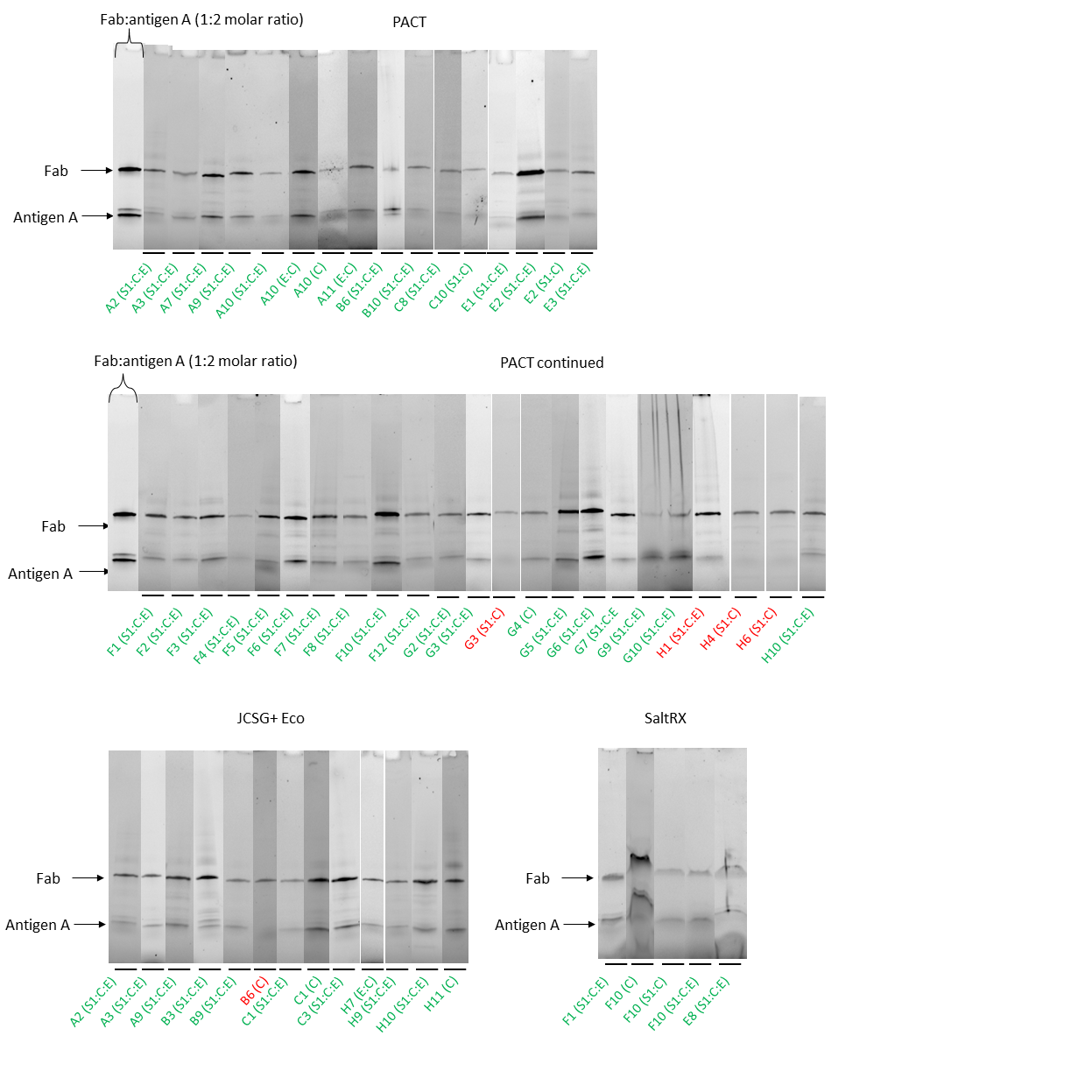

**Supplementary Figure S5. Protein composition analysis for crystals from selected screen conditions of Fab-14836:antigen-A complex variants.** When possible, crystals emerging from JCSG+Eco, PACT and SaltRX screens for Fab 14836:antigen-A complex variants, were picked directly from the 96-well screen trays, washed with precipitant, and analysed by denaturing polyacrylamide gel electrophoresis to assess protein composition. Evaluating the gel images, Fab apo form (red) and Fab:antigen complex (green) were assigned by assessing the absence or presence of antigen-A, respectively, and also by comparing the relative band intensities of Fab and antigen-A. NB: C and E refers to Crystal Kappa and elbow substitutions, respectively.

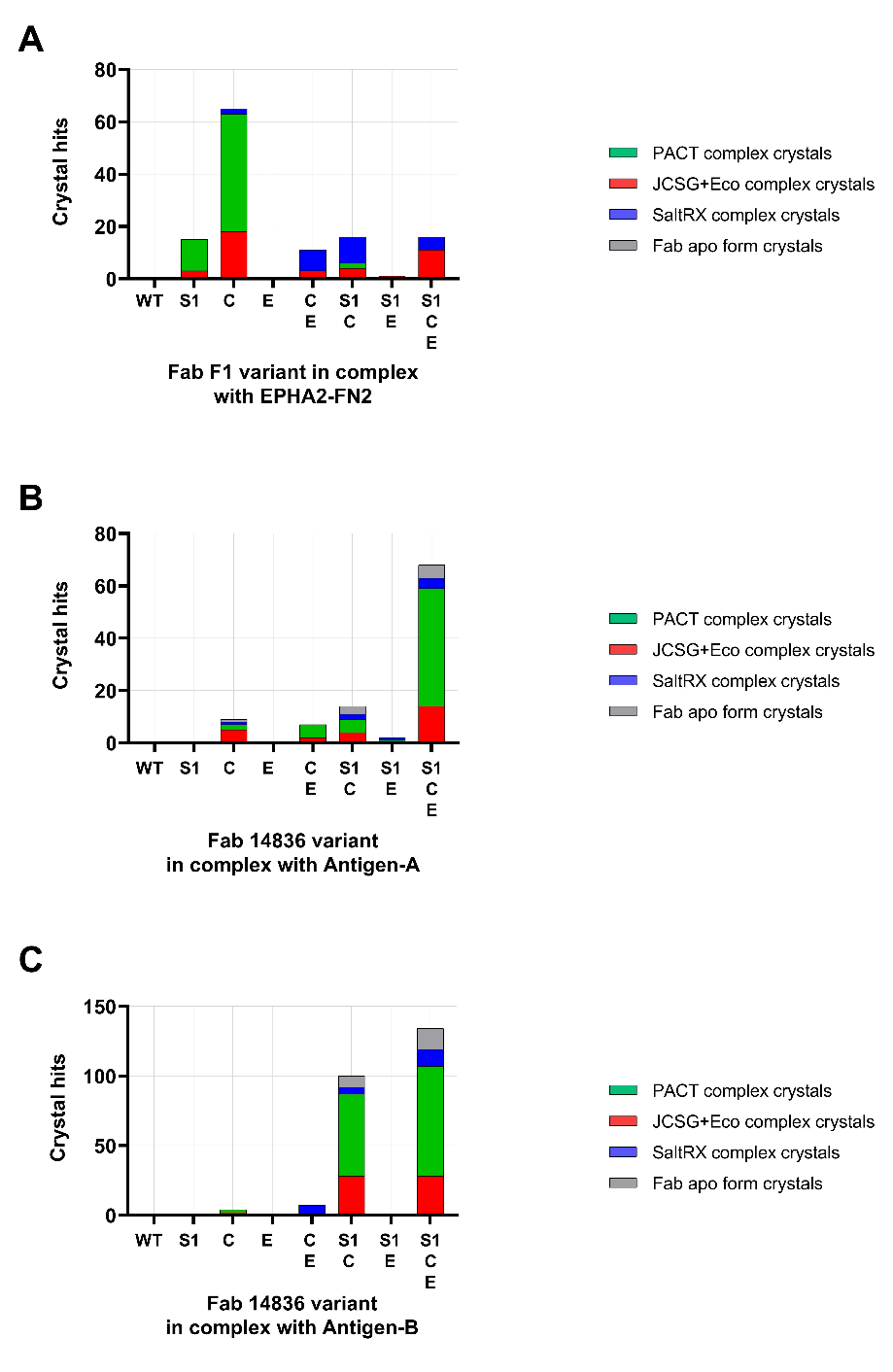

**Supplementary Figure S6. Results of crystallisation screens for Fab variants in complex with antigen.** Results are shown for WT and indicated variants of **(A)** Fab F1:EPHA2-FN2 complex, **(B)** Fab 14386:antigen-A complex, and **(C)** Fab 14386:antigen-B complex. The number of crystal hits (y-axis) obtained for each Fab variant (x-axis) are shown after screening a total of 288 conditions (96 conditions for each of the three indicated screens). The grey bars indicate crystal of apo Fab, as determined by comparing crystal morphology (Fab apo crystallisation drop set up in parallel during the screening - see Supplementary EXCEL doc), and, in the case of Fab 14836:antigen-A, verified where possible by SDS-PAGE (Supplementary Figure S5).

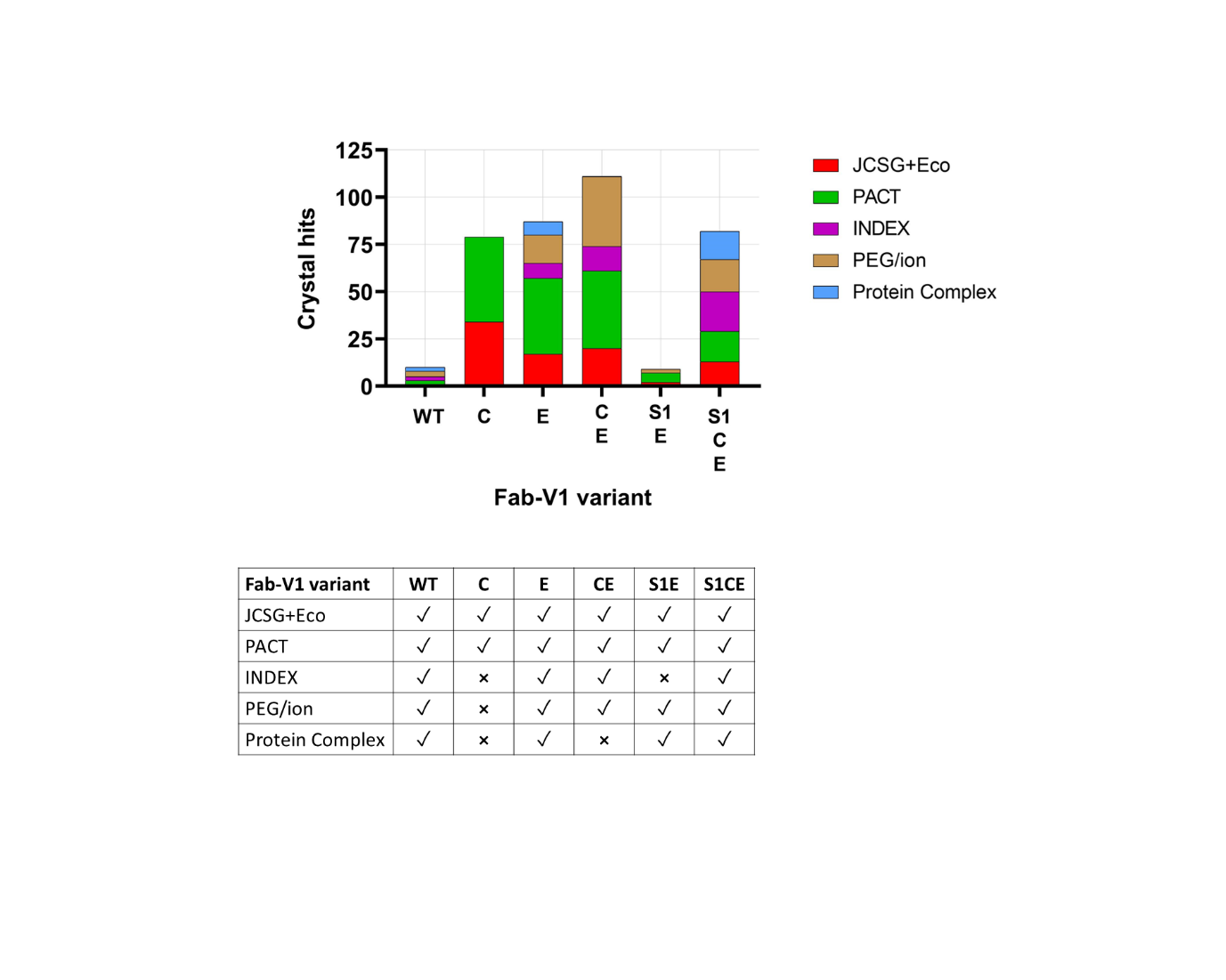

**Supplementary Figure S7. Results of crystallisation screening for selected Fab V1 variants in complex with VHH domain.** The number of crystal hits (y-axis) obtained for each Fab variant (x-axis) are shown after screening a total of 480 conditions (96 conditions for each of the five indicated screens). (Inset Table) details the screens used for each variant.

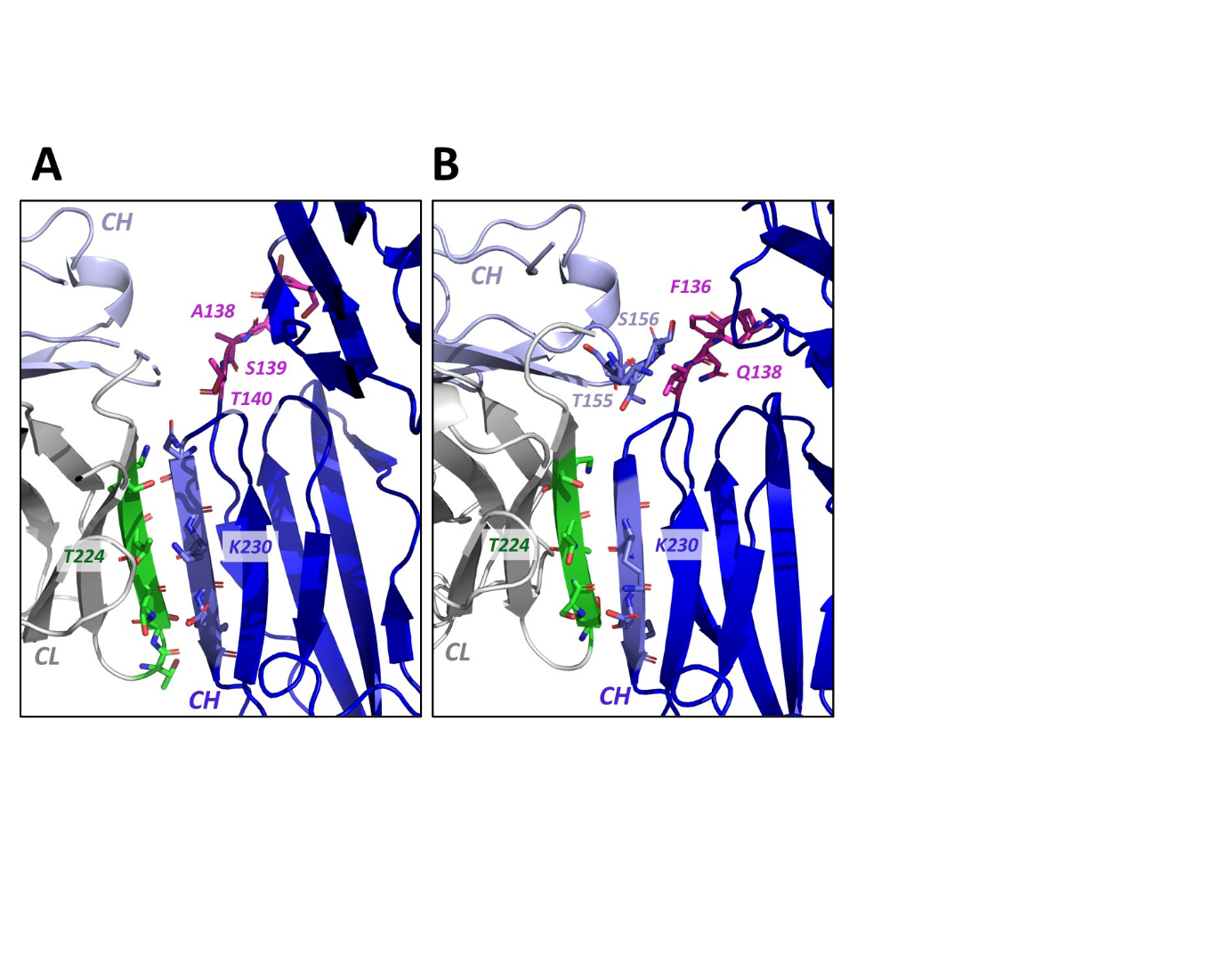

**Supplementary Figure S8. Fab-V1:VHH complex structure Crystal Kappa and elbow substitution crystal lattice packing region.** In **(A)** Fab^C^-V1:VHH, and **(B)** Fab^S1CE^-V1:VHH complex structures, the Crystal Kappa (green) from the principle Fab LC (grey) in the CL domain forms a β-sheet packing interaction with the neighbouring Fab (blue) CH domain. Within the Crystal Kappa packing region, residues T224 and K230 (of the Fab CL domain and packing Fab CH domain, respectively) are labelled for reference. Residues in the WT elbow region (S136, S137, A138, S139, T140) in the Fab^C^-V1:VHH structure do not interact with residues in the neighbouring Fab (light blue) CH domain, which mostly remain unresolved from the electron density - shown in **(A)**. In contrast, hydrogen bond and Van der Waals interactions are formed between the elbow substitution region residues F136, N137, Q138, I139, and residues T155 and S156 in the packing Fab CH domain, in the Fab^S1CE^-V1:VHH complex structure - shown in **(B)**. NB: For more information on crystal lattice packing interactions for these complexes, see Supplementary Table S4.

**SUPPLEMENTARY MATERIAL REFERENCES**

[1]	V. Sobolev, E. Eyal, S. Gerzon, V. Potapov, M. Babor, J. Prilusky, M. Edelman, SPACE: A suite of tools for protein structure prediction and analysis based on complementarity and environment, Nucleic Acids Res. 33 (2005). https://doi.org/10.1093/nar/gki398.

[2]	P. Emsley, B. Lohkamp, W.G. Scott, K. Cowtan, Features and development of Coot, Acta Crystallogr D Biol Crystallogr. 66 (2010) 486–501. https://doi.org/10.1107/S0907444910007493.

[3]	M.-P. Lefranc, C. Pommié, M. Ruiz, V. Giudicelli, E. Foulquier, L. Truong, V. Thouvenin-Contet, G. Lefranc, IMGT unique numbering for immunoglobulin and T cell receptor variable domains and Ig superfamily V-like domains, n.d. http://imgt.cines.fr.
